# Morphogenesis guided by 3D patterning of growth factors in biological matrices

**DOI:** 10.1101/828947

**Authors:** Nicolas Broguiere, Ines Lüchtefeld, Lucca Traschel, Dmitry Mazunin, Jeffrey Bode, Matthias P. Lutolf, Marcy Zenobi-Wong

**Affiliations:** Tissue Engineering and Biofabrication Laboratory, Department of Health Sciences & Technology, ETH Zürich, Zürich, Switzerland; Laboratory of Stem Cell Bioengineering, Institute of Bioengineering, School of Life Sciences and School of Engineering, École Polytechnique Fédérale de Lausanne, Lausanne, Switzerland; Laboratorium für Organische Chemie, Department of Chemistry and Applied Biosciences, ETH Zürich, Zürich, Switzerland

**Author notes:** equal contributions (Ines Lüchtefeld, Lucca Traschel, Dmitry Mazunin).

## Abstract

Three-dimensional (3D) control over the placement of bioactive cues is fundamental to understand cell guidance and develop engineered tissues. Two-photon patterning (2PP) provides such placement at micro- to millimeter scale, but non-specific interactions between proteins and functionalized extracellular matrices (ECMs) restrict its use. Here we report a 2PP system based on non-fouling hydrophilic photocages and Sortase A-based enzymatic coupling offering unprecedented orthogonality and signal-to-noise ratio in both inert hydrogels and complex mammalian matrices. Improved photocaged peptide synthesis, and protein functionalization protocols with broad applicability are introduced. Importantly, the method enables 2PP in a single step and in the presence of fragile biomolecules and cells. As a corollary, we demonstrate the guidance of axons through 3D-patterned nerve growth factor (NGF) within brain-mimetic ECMs. Our approach allows for the interrogation of the role of complex signaling molecules in 3D matrices, thus helping to better understand biological guidance in tissue development and regeneration.

## Introduction

Fluorescent proteins heralded in a paradigm shift in the biosciences in the 1990’s, enabling for the first time the visualization of biological processes in living specimens. The development of a wealth of light-based biosensors followed, notably calcium reporters^1,2^, facilitating the observation of complex processes *in vivo*. A second revolution has been unfolding in the past 15 years, with light being used to control living systems rather than monitoring them^3–9^. In the fields of biomaterials and tissue engineering, this concept has been adopted for controlling, in space and time, the display of extracellular signaling cues to living cells embedded in 3D gels^10^. This technique, coined two-photon patterning (2PP), has opened up exciting perspectives for the *in situ* manipulation of mammalian cells and, in particular, the study of biological guidance^11^. Indeed, 2PP and related methods such as two-photon polymerization^12^ and two-photon ablation^13–15^, have become increasingly useful as biofabrication tools, enabling a potentially unique control over cell and tissue organization at the micro- to millimeter scale^16^.

Because of their modularity and low non-specific interactions, synthetic bioactive hydrogels have been a major focus of 2PP, culminating in the recent ability to reversibly pattern an active growth factor in the presence of living cells^17–21^ in polyethylene glycol (PEG) hydrogels. Nevertheless, despite increasingly sophisticated coupling strategies based on click chemistry and traditional photocages, it has not been possible to utilize 2PP for growth factor-guided tissue morphogenesis, a much sought-after application of this approach. We hypothesized that achieving this milestone would require orthogonal 2PP protocols that are fully compatible with native ECMs, which are most commonly used in fundamental biology studies involving primary cultures or derivation of tissues from self-organizing stem cells. These matrices are however chemically complex, which makes them prone to high levels of signal background from non-specific interactions. Complex biological guidance cues, such as growth factors, are also more prone than simple peptides to adsorption onto traditional photocages, due to their hydrophobicity. Improvements in 2PP patterning specificity, both in terms of physical interactions and of coupling specificity, are therefore crucial. This would ideally be achieved using one-step processes that avoid lengthy incubations and could thus maintain the phenotype and potency of fragile cell types that are currently not photo-modulatable via 2PP.

We therefore devised a 2PP workflow geared towards improved specificity (**Fig. 1**). First, we developed protocols to selectively cage selected amino groups on synthetic peptides with two-photon labile protecting groups. Those were used to cage substrate peptides for Sortase A (SA), a bacterial ligase that provides excellent crosslinking kinetics and outstanding orthogonality to eukaryotic systems^22–25^. We then demonstrated 2PP in inert alginate hydrogels, as well as in an array of naturally derived matrices that are widely used in cell culture and representative of the various mammalian extracellular matrices. Using a hydrophilic photocage was essential to avoid non-specific fouling. Our scheme enables one-step processes, which are not only highly advantageous for *in vitro* 2PP in the presence of cells, but also open future horizons for 2PP, such as the development of *in situ* patterning protocols directly in tissues *in vivo* or *ex vivo*. Furthermore, most labs use standard two-photon microscopes for their patterning and are limited to simple extruded 2D shapes, which we addressed with a new open-source library for advanced 2PP on standard commercial multiphoton microscopes. Finally, we succeeded in guiding axons along defined 3D patterns of nerve growth factor (NGF) in brain ECM-mimetic matrices. These results demonstrate the unprecedented potential of our approach for the 2PP-based induction of tissue morphogenesis.

**Figure 1.**
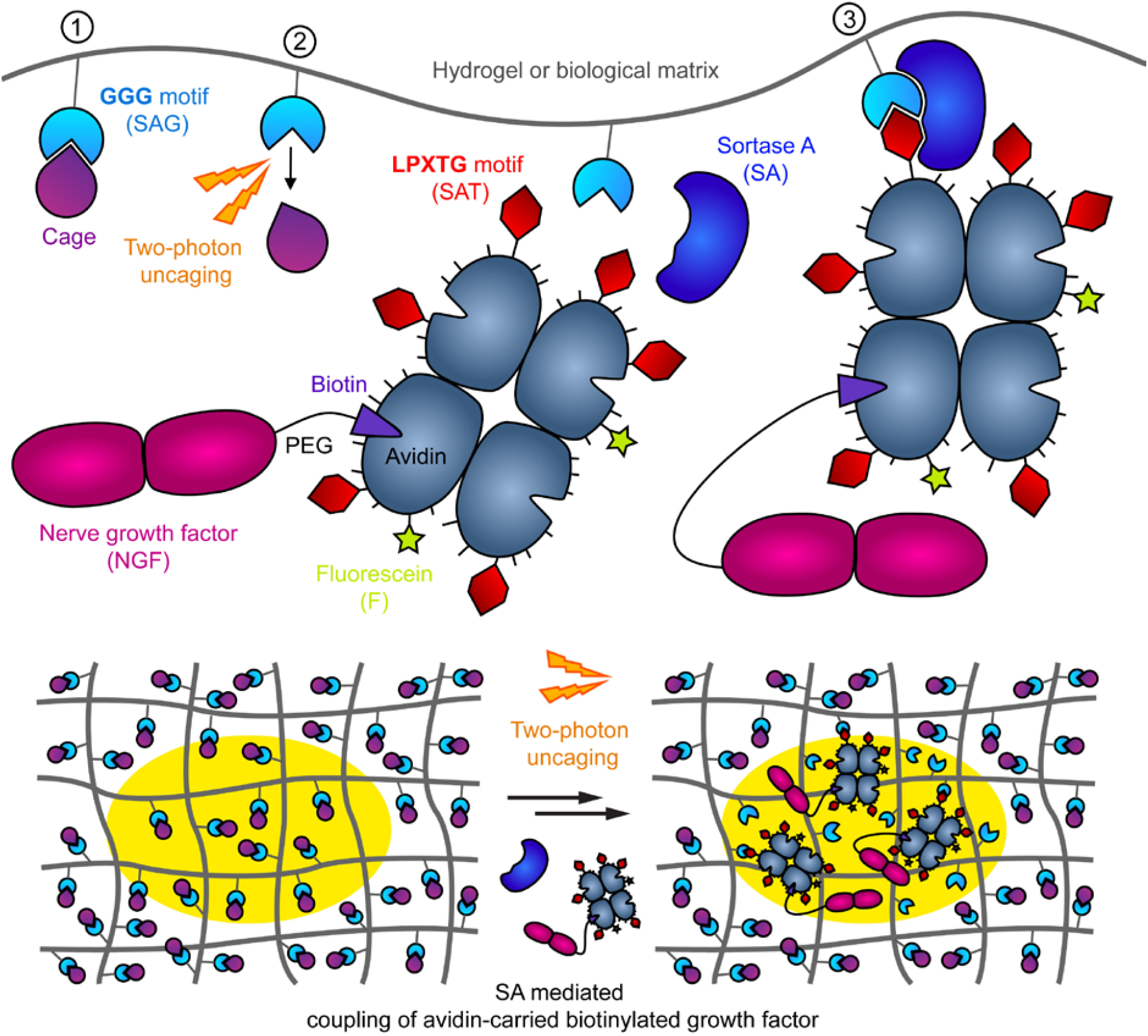
Summary of the 2PP process. In the first step, a hydrogel which harbors caged sortase A (SA) glycine donor peptides (SAG) (1) is formed in the presence of cells and other components needed for the patterning process (i.e. an avidin carrier modified with a SA threonine donor peptide and fluorescein (avidin-SAT-F), SA and biotinylated biological cues). After gel formation (typically within minutes), two-photon uncaging is performed in user-defined positions (2). SA-mediated ligation then anchors the biotinylated biological cue (here, NGF) in the exposed areas (3) through avidin-SAT-F. Note that the growth factor is minimally modified and can be readily exchanged for other biotinylated bioactive cues, that two-photon excitation provides 3D patterning possibilities, and that the resulting patterns can be monitored using the fluorescent tag on the avidin. Bold letters indicate the motifs recognized by SA, in amino acid one-letter code.

## Results and discussion

### Flexible synthesis of two-photon active caged peptides

We synthesized peptides inactivated by various photocages. We studied the traditional 2-(2-nitrophenyl) propoxycarbonyl (NPPOC)^26^, which is widely used in the synthesis of DNA arrays and has been used to cage amino-acids^27^, but is known to be relatively insensitive to two-photon uncaging in the absence of a sensitizer^28^, as other cages derived from o-nitrophenyl. We then studied 6-bromo 7-hydroxycoumarin (BHC), the first described cage with high two-photon uncaging cross-section^29,30^, as well as a newer variant with improved stability, 7-diethylaminocoumarin (DEAC)^31,32^. Finally, as we were concerned with the hydrophobicity of these traditional cages, we studied a more recently described variant of DEAC, 7-dicarboxymethylaminocoumarin (DCMAC)^33^, which has extremely favorable water solubility and two-photon uncaging properties^34^. DCMAC has only been sparingly used, despite of its interesting features, most likely due to the lack of a high yielding synthesis. This led us to develop a new procedure, performing the reaction solvent-free in the presence of a radical inhibitor, leading to 80-90% yield in the critical first step of the synthesis.

Photocages could be readily added on the N-terminus of solid phase-supported peptides after 4-nitrophenyl chloroformate activation. Conveniently, the resulting carbamate linkages resisted standard peptide cleavage conditions, notably including concentrated aqueous trifluoroacetic acid and thiols. This method was used to cage SAG peptides with DCMAC (**Fig. 2a**) and other cages (**Fig. S1–3**). Interestingly, the protocol can be readily combined with base-insensitive orthogonal protecting groups such as Ddiv, to enable the selective caging of any side chain amino group. As an example, we caged on its lysine side chain the peptidic substrate (**FKGG-ERCG**) of the human transglutaminase activated factor XIII (TG), which yielded functional two-photon activatable TG substrates (**Fig. S4a-b**). The versatility of the procedure is an attractive feature for the generation of both hydrophobic and hydrophilic two-photon activatable bioactive peptides.

**Figure 2.**
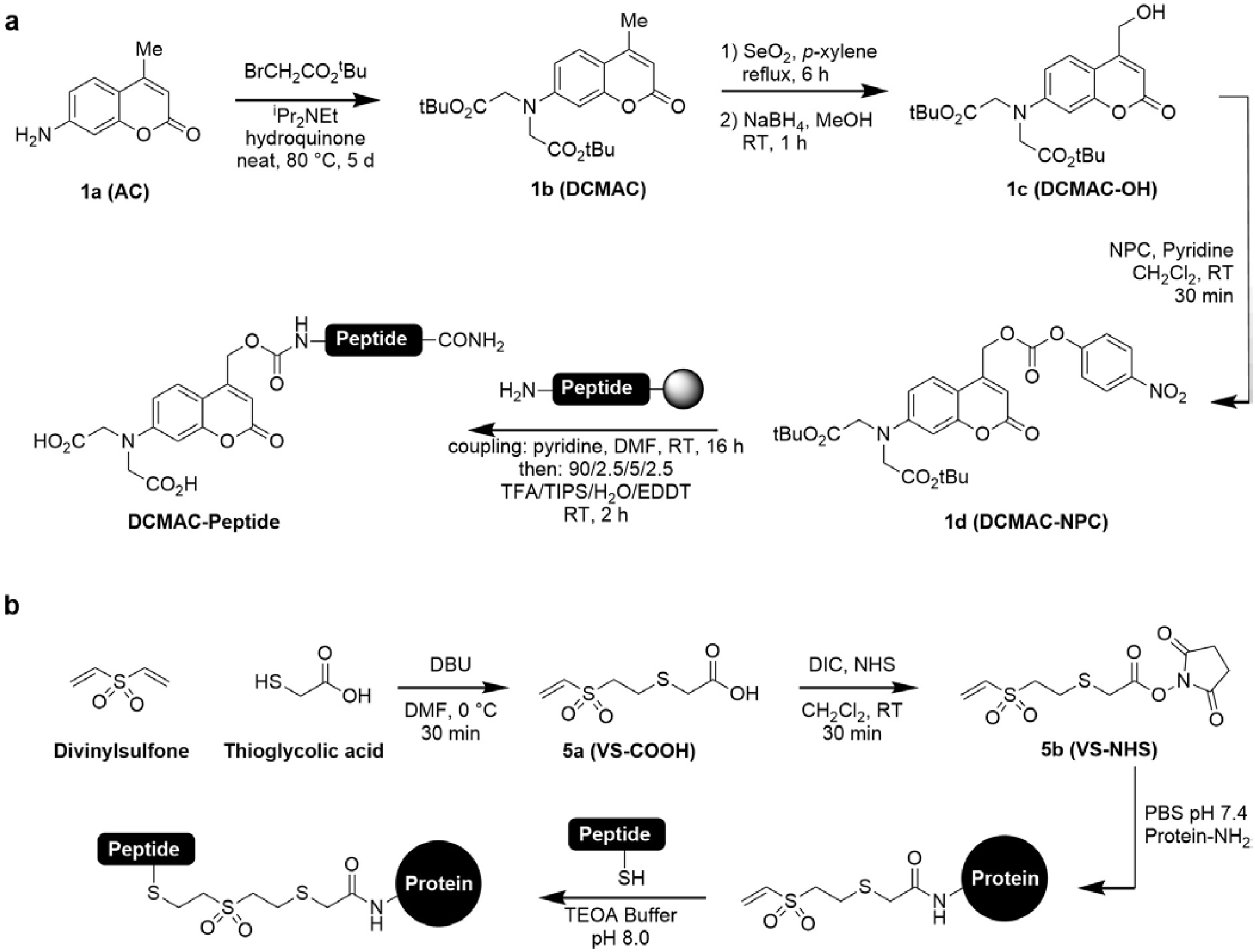
Synthesis of (**a**) the hydrophilic cage DCMAC and caged peptides, and (**b**) the heterobifunctional crosslinker VS-NHS and stable protein-peptide conjugates. Abbreviations: AC: 7-amino 4-methylcoumarin, DCMAC: 7-dicarboxymethylaminocoumarin (and by extension the 4-methyl and ^t^Bu protected derivatives), NPC: 4-nitrophenyl chloroformate and 4-nitrophenyl carbonate ester, TFA: trifluoroacetic acid, TIPS: triisopropylsilane, EDDT: 2,2’-(ethylenedioxy)diethanethiol, DBU: 1,8-Diazabicyclo[5.4.0]undec-7-ene, TEOA: triethanolamine, PBS: phosphate buffered saline, VS: vinylsulfone, NHS: N-hydroxysuccinimide, DIC: N,N’-diisopropylcarbodiimide.

### Development of a facile strategy to generate photo-patternable proteins

Developing new tagged proteins is a time-consuming and expensive process, especially when it involves cloning and recombinant expression. Biotinylated proteins on the other hand are already a standard, and most growth factors are readily available in a biotinylated format. We therefore devised a flexible 2PP strategy based on functionalized avidin, that could act as a carrier by binding biotinylated growth factors.

In order to decorate fluorescein-tagged avidin with thiol-bearing SAT peptides, we investigated several classical amine-thiol heterodifunctional cross-linkers with limited success. A few problems encountered were low substitution efficiency with maleimide-PEG-NHS 3.4 kDa (likely due to the large polymer chain crowding the NHS environment), lysis of linkages created by Traut’s reagent in the presence of thiols at physiological pH, and retro-Michael addition from maleimide linkers, all known issues^35,36^.

This motivated the synthesis of a short and highly reactive heterodifunctional cross-linker, vinylsulfone-(S)-glycolic acid-(N)-hydroxysuccinimide ester (VS-NHS, 5b), that is straightforward to synthesize (Fig. 2b) and gives highly stable^37^ amide and thioether linkages (Fig. 2b). Remarkably, based on these properties, VS-NHS would be an advantageous alternative to the commonly used cross-linker succinimidyl 4-(N-maleimidomethyl) cyclohexane-1-carboxylate (SMCC) for the construction of protein conjugates, avoiding altogether hydrophobic spacers and retro-Michael addition issues. This route was used to prepare highly substituted avidin, with 6 SAT peptides per avidin, as demonstrated by mass spectrometry (**Fig. S5a-b**). The binding of biotinylated NGF, used as a model growth factor, to the substituted avidin-SAT-F was confirmed by gel permeation chromatography (Fig. S5c-f).

### Non-fouling enabled by hydrophilic photocages is critical to 2PP performance

Next, we performed a side-by-side comparison of photocages to see how the choice of the cage influences the brightness and specificity of 2PP using SA-mediated patterning of avidin-SAT-F as model system (**Fig. 3a-c**). To minimize background signal, we used inert non-fouling alginate hydrogels as a blank canvas. Optimal laser intensities were determined for each cage. Whereas low laser power failed to uncage, too high power was equally detrimental, causing photodamage of ligands. The optimal power for one-scan uncaging was 10-40% laser power at a frame rate of 0.6 Hz, depending on the cage, similar to that used for two-photon imaging. Patterns obtained with the hydrophilic two-photon active cage DCMAC were highly specific, whereas traditional cages only enabled faint patterns that were barely above background.

**Figure 3.**
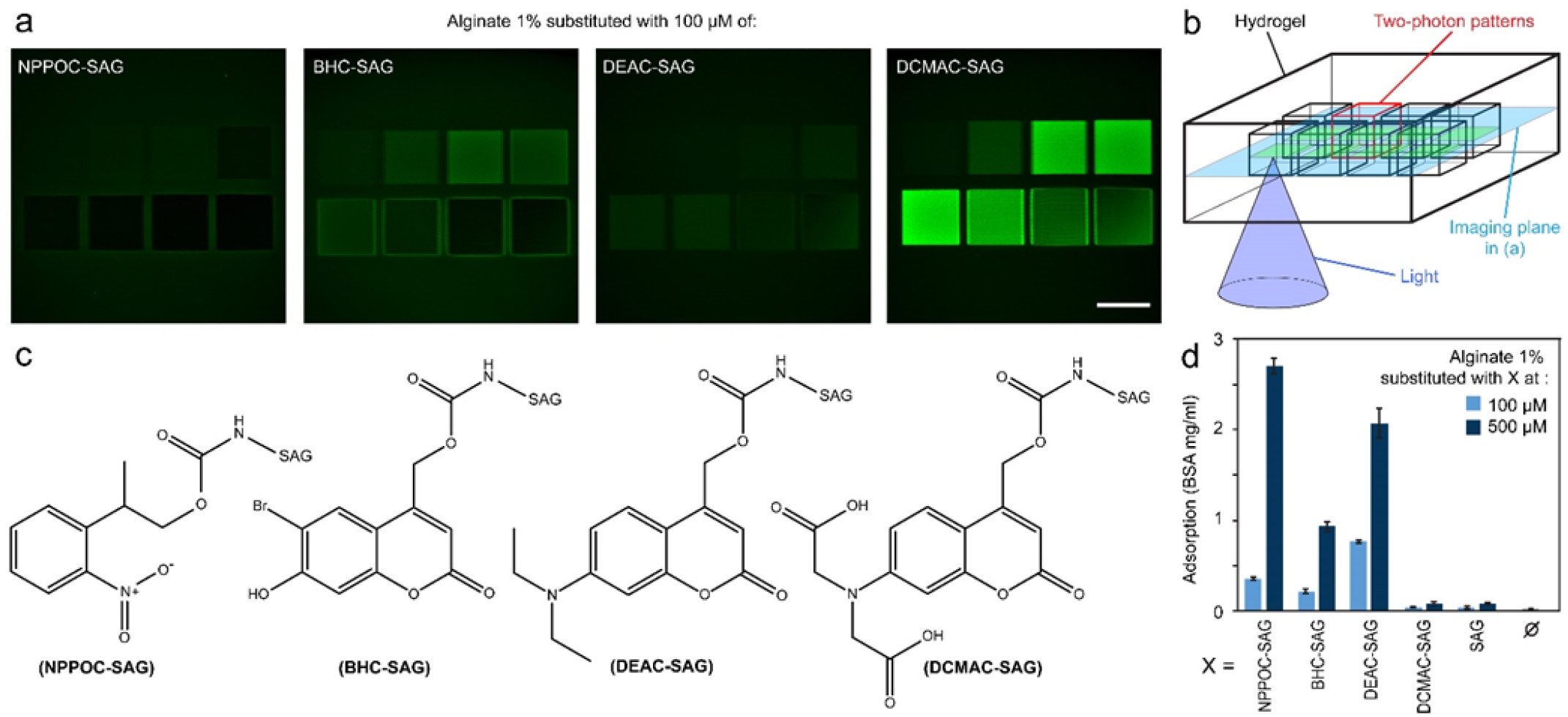
Comparison of photocages. (**a**) Two-photon patterns of avidin in alginate obtained with different cages. Scale bar: 100 μm. Coupling is done with Avidin-SAT-F 0.35 mg/ml, SA 4 μM, CaCl_2_ 10 mM. Uncaging is done at 770 nm with a fs-pulsed Mai Tai laser at laser powers of 2.5, 5, 10, 20% (top row) and 30, 40, 60, 80 % (bottom row). Confocal imaging of the fluorescein reporter mid-depth of the cubes was acquired with quantitative photon counting detectors and is displayed with identical brightness and true background values. (**b**) Schematic of the experimental setup: the hydrogel is functionalized with avidin on a 3D array of 100 μm cubes. (**c**) Chemical structures of the cages tested on SAG peptides. (**d**) Quantification of the non-specific adsorption of rhodamine-tagged bovine serum albumin (BSA) onto hydrogels substituted with caged peptides and controls. The BSA was incubated at 2 mg/ml for 100 min, and fluorescence readout was done after 48 h of washing. Error bars: SD, n=4.

Hydrophobic surfaces are known to induce strong adsorption of proteins^38^, and we therefore hypothesized that the poor outcomes with traditional cages, which are quite hydrophobic, might be primarily due to protein adsorption, increasing the background close to the surface and depleting the gel of free mobile proteins available for specific couplings in the core. We tested this hypothesis by quantifying the adsorption of bovine serum albumin (BSA), the serum protein most typically used for adsorption studies, to alginate hydrogels modified with various moieties (Fig. 3d). A rhodamine reporter was conjugated to the BSA for sensitive fluorescent readout using red-light that does not interfere with the photocages. We found that unmodified alginate retains little adsorbed protein (25±5 μg/ml, SD n=8), and alginate substituted with unmodified SAG or DCMAC-SAG at a concentration of 100 μM, typical for 2PP, did not retain significantly more BSA (p>0.75). Other cages at the same concentration retained from 5 to 20 times more protein (p<1e-7). Higher concentrations exacerbated the difference even further, with 10 to 40 times more adsorption on traditional cages than on DCMAC. Non-specific adsorption is therefore an essential property that should be optimized for high quality 2PP.

### High specificity enables 2PP in complex biological matrices and one-step processes

After identifying the superior performance of DCMAC, we next generated an array of DCMAC-caged SAG peptides bearing a variety of handles for easy incorporation in all common synthetic and biological matrices (**Fig. 4a**). Equipped with these tools, we demonstrated 2PP in a collection of mammalian matrices, representative of the ECM of connective (collagen), epithelial (Matrigel), central nervous system (hyaluronan^39^), and regenerative (fibrin) tissue (Fig. 4b). Remarkably, optimal uncaging could be achieved in a single scan. This enables relatively fast pattern formation in around 1.5 min for patterns of 500×200×200 μm. Higher exposures resulted in damage to the hydrogel and peptide handles rather than increased uncaging, as evidenced by a reduction in the coupling efficiency (e.g. Fig. 4b with 40% laser power and 4 scans in fibrin and hyaluronan).

**Figure 4.**
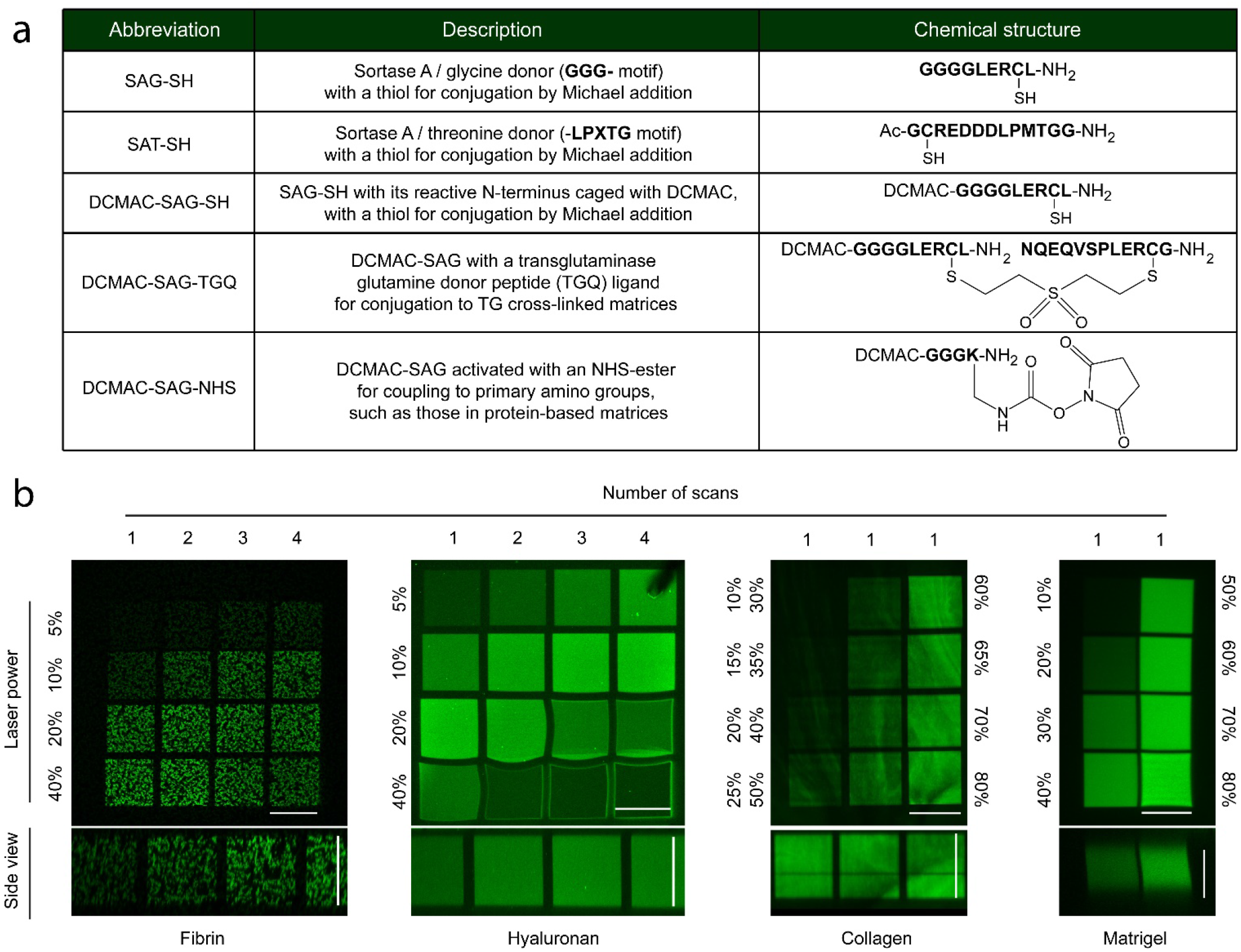
2PP in biological matrices. (**a**) Peptides used as substrates for enzymatic couplings, and caged derivatives used to functionalize various gel backbones. Characters in bold are amino acids in one-letter code. Critical functional groups on the side chains of amino acids used for couplings are highlighted in explicit chemical notations. -NH2 indicates amidated C-terminus, and Ac-indicates acetylated N-terminus. (**b**) 2PP of avidin-SAT-F in the main mammalian extracellular matrices, uncaging each test cube with a different laser power or scan number. Confocal fluorescence images of the fluorescein reporter were acquired with quantitative photon-counting detectors, with backgrounds not subtracted in order to display true non-specific adsorption outside of the patterns. The images in collagen are average intensity projections over the whole cube thickness to mitigate the inhomogeneities due to collagen fibers, other images are single planes. Patterns in fibrin and hyaluronan were produced on a Leica SP8 inverted with a 25x water immersion objective, collagen and Matrigel on an SP5 inverted with a 20x water immersion objective. Scale bars: 100 μm.

The enzymatic reactions used for hydrogel cross-linking (TG, thrombin) and 2PP couplings (SA) are fully orthogonal and compatible with physiological conditions, bypassing the need for serial incubations, which were necessary in previous photopatterning systems^18,20,21,30,40,41^. This advance is key to reduce the handling time and the exposure of 3D encapsulated cells to soluble bioactive cues, and could also open the way to *in situ* patterning after delivery into a tissue defect *in vivo*, an application that is of utmost interest for aligned tissue reconstruction. For example, *in vivo* 2PP might be used to create guidance channels in damaged neural tissue that match the surrounding structures, or to alter guidance in developing embryos in order to study the mechanisms of morphogenesis. We tested the feasibility of this one-step process *in vitro* by including the modified avidin, growth factor, as well as gel-cross-linking and 2PP-coupling enzymes in the same gelling mix as the biopolymer. Gelation occurred within minutes, which enables immediate photopatterning and transfer to cell culture. This incubation-free one-step process worked equally well as a classical two-step procedure (Fig. S4c).

### Patterning complex gradients and 3D shapes on standard microscopes

While two-photon polymerization instruments are common in microfabrication facilities, and advanced 2PP setups for biological applications have been built by a handful of bioengineering labs^42,43^, most biologists can only rely on standard two-photon microscopes for their two-photon uncaging. Scanning a region of interest (ROI) over a 3D stack or continuously changing the laser power enables the formation of extruded 2D-shapes or simple gradients on such instruments, but more complex gradients or truly 3D shapes are out of reach. In an effort to make advanced 2PP available to biologists using existing infrastructure, we developed an open-source Matlab library that automatically generates instructions for standard commercial Leica two-photon microscopes (**Fig. 5**).

**Figure 5.**
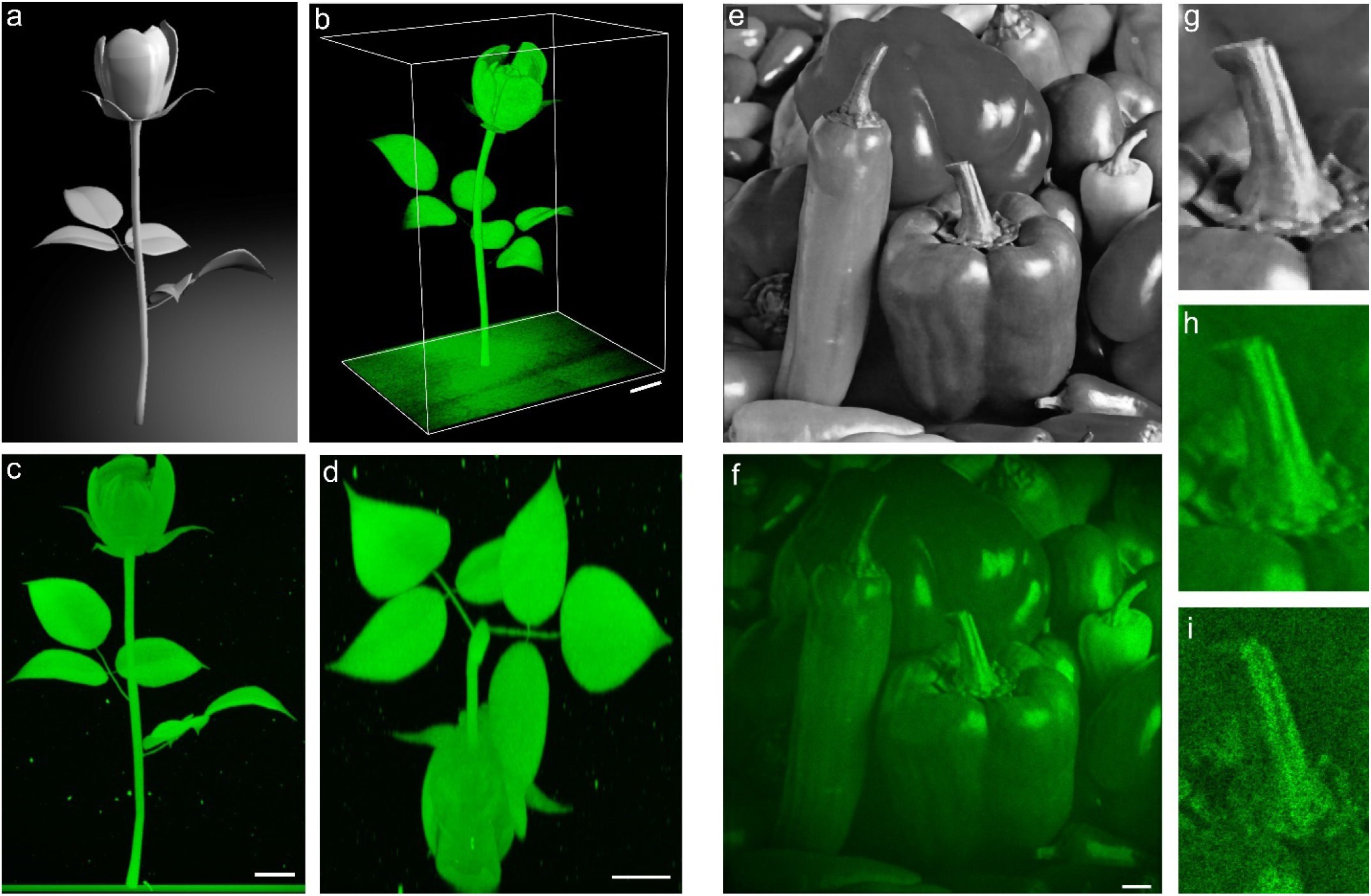
2PP of complex gradients and 3D shapes using a standard commercial two-photon microscope. (**a**) 3D model and (**b**) 2PP of avidin-SAT-F in a transglutaminase cross-linked hyaluronan hydrogel using a one-step process, as seen with two-photon imaging of the fluorescein tag (Fiji 3D viewer surface view). (**c, d**) Maximum intensity projections in orthogonal planes. (**e**) Grey-scale image exhibiting complex gradients, and (**f**) reproduction as an avidin-SAT-F pattern in a collagen matrix (average intensity projection of a 200 μm thick pattern, invariant in the z direction). (**g, h**) Close-ups of the reference and pattern. (**i**) Single-plane view, with the collagen microstructure visible in the absence of averaging. Scale bars: 50 μm.

In order to provide some robustness to various machine setups and facilitate setting the parameters that are kept constant (e.g. scanning speed, objective, filter positions), our scripts use a template exported from the microscope. Complex instructions are generated based on this template, by adjusting the laser power, z-positions, and scan areas. The scripts were tested on different instruments, including a Leica SP8 inverted with a Spectra Physics Mai Tai laser (Fig. 5a-d) and an SP5 upright with a Chameleon laser (Fig. 5e-i).

We then ensured the patterning instructions can be given in standard open formats: complex 3D shapes are input as triangular mesh data in the stereolithography (.stl) format, that can be exported/converted from most 3D modeling software, including open-source software such as Blender and Meshlab. The scripts slice the mesh to create a series of closed paths, which are then converted to a series of regions of interests associated with a z-position, selectively exposed during 3D scan using standard built-in electro-optical modulators (EOM) as a switch. Complex gradients are input as pixel data, typically tagged image format (.tif) files, together with calibration curves of pattern intensity vs uncaging conditions and laser power vs hardware filter position. Masks associated to different intensity levels in the reference image are then serially exposed to the appropriate laser powers, as defined by the reference curves, for accurate patterning.

Using this library, we could reproduce fine intricate 3D models (Fig. 5a-d) as well as complex gradients (Fig. 5e-i) as micropatterns of functionalized avidin. Of note, the fidelity of the patterning process depends on the optical system rather than on our chemical or software tools. In particular, the patterning resolution is defined by the size of the point spread function, which is linked to the choice of the objective and is well known from standard two-photon microscopy. We believe our algorithms will highly facilitate the use of two-photon fabrication/functionalization techniques in life science research, turning commonly available commercial microscopes into flexible two-photon fabrication instruments.

### Axons follow nerve growth factor patterns in a brain-mimetic matrix

The development of the complex architecture of the nervous system arises from patterns of morphogens which guide axonal outgrowth. Models recapitulating this type of biological guidance would ideally use 3D rather than 2D cultures, for physiologically relevant growth cone cytoskeletal organization. They would also use biological or biomimetic background matrices providing a relevant microenvironment.

To implement this concept, we chose sensory neurons from dorsal root ganglia (DRG) explants of E9-10 chick embryos as a model system, well known for its response to NGF affecting neuron survival and axonal growth^44^. Since biological matrices are likely to interfere with the response to the growth factor, we first compared the axonal growth from DRGs with or without soluble NGF in the medium, in an array of biological matrices (**Fig. S6**). Ideally, a good matrix for precise guidance would support axonal growth in the presence of the growth factor and inhibit it otherwise. Matrigel, collagen and fibrin supported extensive glial and axonal outgrowth both in the presence and absence of NGF, consistently with previous findings^44–46^, which limits their usefulness here. Hyaluronan gels, on the other hand, were almost entirely resistant to DRG axonal growth in the absence of NGF, but moderately supportive in the presence of soluble NGF, which is coherent with previous reports that soluble hyaluronan is inhibitory for axonal growth from DRGs^47^ and notably different to the fast neurite outgrowth from central neurons in these gels^39^. Axonal growth was found to be much enhanced in the presence of immobilized growth factor (**Fig. S7c**), a rather surprising finding even though there are prior reports of moderately increased axonal growth upon immobilization of NGF^48,49^. Hyaluronan was therefore chosen as an ideal background.

Micropatterns of NGF were formed in apposition with freshly encapsulated DRGs. When NGF was immobilized in large regions (200×500×500 μm^3^), axons extended at a fast rate of ≈100 μm/day and stayed confined in the area, which was particularly striking as axons collectively took sharp 90° turns when reaching pattern edges (**Fig. 6a-e**). This demonstrates the robustness of the approach, and gives a bioengineered model of how anchored morphogens can strictly restrict cell processes to the tissues to which they belong.

**Figure 6.**
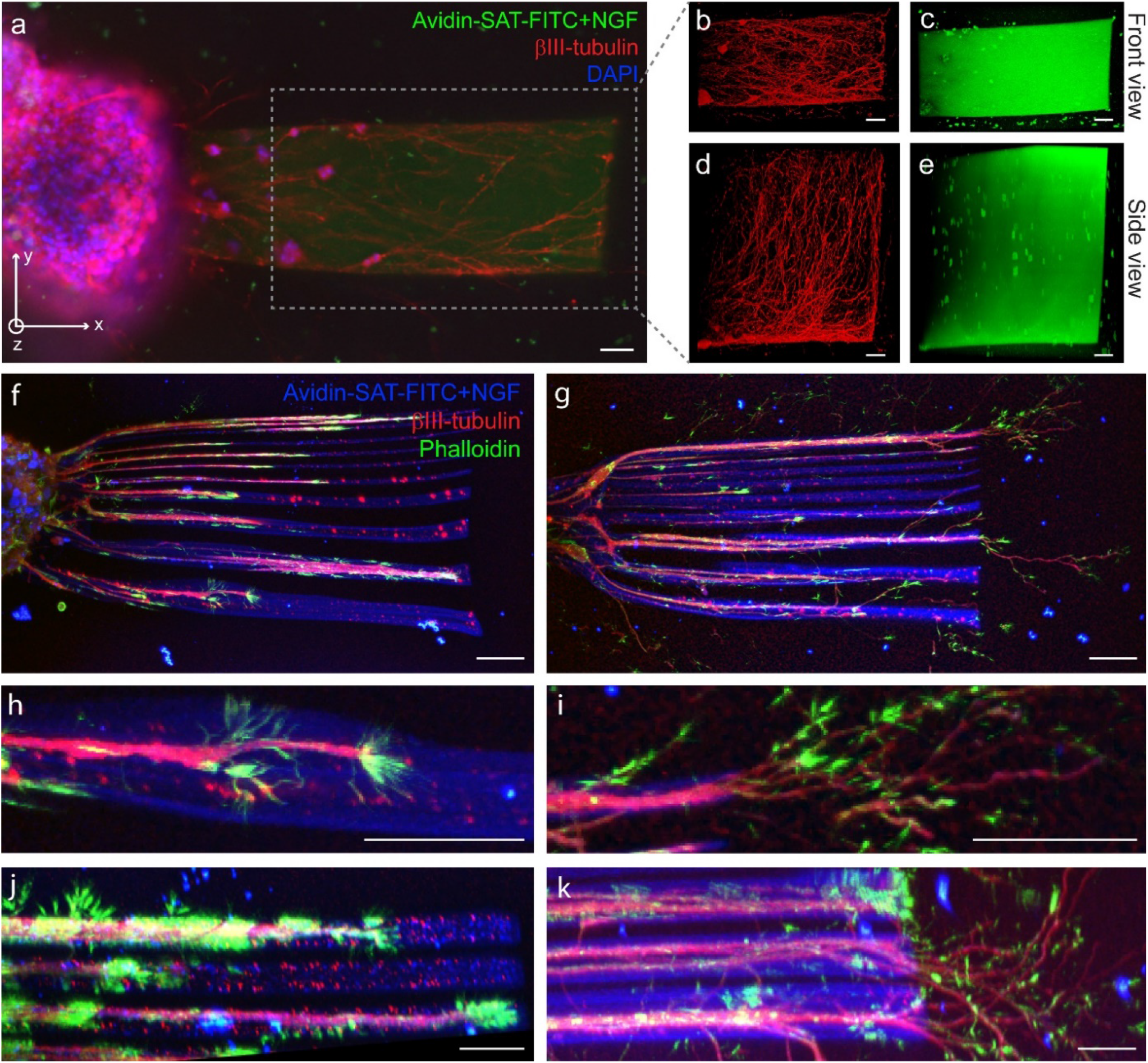
Axonal guidance from chick dorsal root ganglia (DRG) into transglutaminase cross-linked hyaluronan matrix patterned with nerve growth factor (NGF) (**a-e**) Representative images of axon confinement within a large rectangular NGF-positive region (200×400×400 μm^3^) after 10 days in vitro. (**a**) Single plane widefield fluorescence imaging showing the DRG as a DAPI-positive structure on the left, the pattern as a fluorescein-tagged rectangle on the right, and βIII-tubulin-positive axons mostly confined to the patterned area. (**b-e**) Maximal intensity projections from confocal imaging data of the pattern area in (**a**), with (**b-c**) projected along z and (**d-e**) along the y axis. (**f-k**) Representative images of axon guidance with microchannels of NGF (2 to 40 μm wide from top to bottom), after 2 days of culture. Maximum intensity projections from confocal z-stacks. (**h-i**) and (**j-k**) are close-ups from (**f-g**) in front and side view respectively. Images are adjusted for brightness/contrast/gamma for clarity. Scale bars: 50 μm.

In a classical understanding of NGF guidance, chemotaxis follows gradients of the soluble growth factor. Nevertheless, other mechanisms might have been missed due to the lack of experimental systems enabling 3D immobilization. We hypothesized that microchannels with dimensions comparable to the size of a growth cone could effectively orient axonal growth, similarly to lines of Schwann cells during nerve regeneration. We therefore patterned microchannels of 2-40 μm width in front of encapsulated DRGs (Fig. 6f,h,j). Interestingly, axons grew as bundles strictly aligned with the microchannels, and upon reaching the end, de-bundled to form an arborization (Fig. 6g,i,k). In order to ensure reproducibility, the DRG guidance results were repeated 3 times, using at least 2 different batches for the synthesis of each compound, with similar results. We also ensured that the guidance is specifically mediated by immobilized NGF by performing mock patterning, replacing biotinylated-NGF with non-biotinylated NGF, which prevented the growth (Fig. S7a-b). The bundles and trees are reminiscent of the behavior of axons that grow along a tract and spread upon reaching their target tissue, and this reproduction in a bioengineered system sheds new light on how the microenvironment can guide these morphogenetic events.

### Conclusions

Our protocol for 2PP based on an orthogonal enzymatic coupling and non-fouling hydrophilic photocages enabled the patterning of avidin-linked biotinylated growth factors in mammalian extracellular matrices. Patterns of NGF in hyaluronan gels successfully guided axonal growth from sensory neurons in 3D. The system therefore introduces new opportunities for studying the mechanisms involved in axonal guidance by NGF signaling in three-dimensional space. Importantly, it also lays the foundation for guidance studies based on other cues, cells, and matrices, as the methods are straightforward to translate to other biotinylated growth factors / morphogens due to the versatility of avidin as a carrier. The low off-target interactions of the optimized building blocks act as a safety net against toxicity, make intermediate washing or blocking steps unnecessary, and limit background from non-specific protein binding.

2PP of sensitive biomolecules in complex matrices being particularly demanding, we were led to develop some chemical tools that might be of interest for use in much broader fields. First, our new heterodifunctional cross-linker, VS-NHS, could be an interesting alternative to the maleimide-NHS SMCC to form protein conjugates with improved linkage stability and reduced hydrophobicity. Our high yielding synthesis of a hydrophilic two-photon active cage could be incorporated in other two-photon patterning strategies, in order to avoid non-specific adsorption when patterning proteins. Our protocols to protect peptides with weakly interactive and highly water soluble two-photon active cages on any amino group could lead to improved substrates to study the signaling events generated by biologically active peptides with precise control over 3D position and time.

Further improvements in 2PP protocols could include multi-color extension of one-step high-specificity 2PP: the development of hydrophilic cages activated at orthogonal wavelengths together with the validation of additional orthogonal enzymes/ligands would be a natural path forward. Finally, this first demonstration of a one-step protocol to pattern growth factors in biological matrices and guide morphogenesis should motivate the development of *in vivo* 2PP, aiming towards in situ micro-aligned guidance of tissue regeneration.

### Code availability

The Matlab library for Free-form 2-photon patterning (F2P2) is available at: https://github.com/nbroguiere/F2P2.

## Acknowledgements

We would like to thank David Fercher for technical assistance with peptide and protein production, and Prof. Esther Stoeckli and Dr. Beat Kunz for help with DRG isolation. We are grateful to Konstantin Schulz-Schönhagen, Michèle Dai, and Jasmin Feike for technical assistance during the preliminary exploration of cage synthesis options. We further acknowledge the assistance from the imaging facilities of the ETH (ScopeM) and EPFL (BIOP), with particular thanks to Dr. Justine Kusch and Thierry Laroche.

## Materials and methods

### General procedures

All chemicals are from Sigma-Aldrich-Merck, and cell culture reagents from ThermoFischer Scientific, unless indicated otherwise. NMR spectra were acquired on a Bruker AVIII400 or similar instruments and are provided in annex. High performance liquid chromatography (HPLC) was performed on C18 columns using Agilent instruments, using Poroshell 120 EC-C18 4.6 x 150 mm, 2.7 μm particle size columns for analytical HPLC, with a gradient from 10 to 90% acetonitrile (ACN) in water over 5 min in the presence of 0.1% trifluoroacetic acid (TFA) at 2 ml/min and 50°C, unlike indicated otherwise. Preparative HPLC were on an Agilent Prep C18, 100 Å pore size, 10 μm particle size, 20 x 150 mm column at a flow rate of 20 ml/min and room temperature, for up to 50 mg of peptide, or the equivalent 50 mm diameter column at a flow rate of 80 ml/min for up to 300 mg of peptide, using gradients of 10 to 90% ACN in water over 40 min in the presence of 0.1% TFA unless indicated otherwise. Gel permeation chromatography (GPC) were on an AdvanceBio SEC 300 Å pore size, 2.7 μm particle size, 7.8 x 300 mm column from Agilent. Liquid chromatography – mass spectroscopy (LCMS) were acquired on a Waters HPLC instrument equipped with electrospray ionization (ESI) and an SQ. detector 2. Matrix-assisted laser desorption and ionization time-of-flight mass spectrometry (MALDI-TOF) on α-cyano-4-hydroxycinnamic acid (HCCA) matrix in the presence of TFA were acquired on a Bruker Autoflex instrument. Peptide sequences in amino-acid one-letter code are highlighted in bold throughout the methods.

### Synthesis of 7-di-tert-butyl-carboxymethylamino 4-methylcoumarin (DCMAC, 1b)

1 g (5.7 mmol) of 7-aminocoumarin (AC, 1) was mixed with 3 ml (17.2 mmol) of ^i^Pr_2_NEt, 3 ml (20 mmol) of tertbutyl bromoacetate (BrAcOtBu), and 100 mg of hydroquinone in a Schlenk tube. The tube was sealed and the slurry was left to react at 80 °C for 5 days in the dark with magnetic stirring. At this point, the slurry was dissolved in dichloromethane (DCM), washed with water and brine, dried over MgSO4, and purified by flash chromatography (EtOAc:Hexane 1:2) with dry loading. The compound (1.91 g, 83%) was recovered as a white crystalline solid after drying under reduced pressure. Rf=0.6 (EtOAc:Hexane 1:1).

### Synthesis of 7-di-tert-butyl-carboxymethylamino 4-hydroxymethylcoumarin (DCMAC-OH, 1c)

2 g (5 mmol) of DCMAC 1b were refluxed under nitrogen for 6h with 1.1 g (10 mmol) of selenium dioxide in 25 ml of p-xylene. Insoluble tars were removed by hot filtration, and the solvent was removed on a rotovap. The compound was resuspended in 40 ml of MeOH, and 190 mg (5 mmol) of NaBH_4_ were added during stirring. After 30 min at room temperature, the pH of the mixture was balanced to 2 by addition of aqueous HCl 1M, and further diluted with water before extracting 4 times with DCM. The combined organic phases were washed with brine, dried over MgSO4, and evaporated onto silica gel for purification by flash chromatography with dry loading (EtOAc:Hexane 1:1). Upon drying, a dense dark yellow solid was obtained, which was resuspended in 10 ml of dimethyl formamide (DMF) and further purified by C18 preparative HPLC in a gradient from 40 to 100 % ACN in water with 0.1% TFA. After lyophilization, 1 g (48%) of the compound was obtained as a light off-white powder. Rf=0.2 (EtOAc:Hexane 1:1).

### Synthesis of (7-di-tert-butyl-carboxymethylamino coumarin 4-yl)methyl (4-nitrophenyl) carbonate (DCMAC-NPC, 1d)

335 mg (0.8 mmol) of DCMAC-OH 1c and 0.3 ml (3.72 mmol) of pyridine were dissolved in 4 ml of dry DCM, and 153 mg (0.8 mmol) of 4-nitrophenyl chloroformate (NPC) pre-dissolved in 2 ml of DCM were added dropwise to this solution while stirring. The activation was monitored by analytical HPLC (40 to 100% ACN in water, with 0.1% TFA), and additional NPC was added if needed. The reaction was completed in around 30 min. The reaction mixture was then dried under reduced pressure, and used in the next step without further purification, since the 4-nitrophenol and pyridinium chloride byproducts do not interfere with the following reaction. Rf=0.6 (EtOAc:Hexane 1:1).

### General procedures for peptide synthesis

Peptides were synthesized with classical Fmoc-protected solid-phase supported synthesis on rink-amide resin, using (1-Cyano-2-ethoxy-2-oxoethylidenaminooxy)dimethylamino-morpholino-carbenium hexafluorophosphate (COMU) as a coupling reagent in the presence of ^i^Pr_2_NEt, and 20% piperidine in DMF for Fmoc deprotection. Peptides were dried under nitrogen flow in the reaction vessel, cleaved and deprotected by incubation for 2 h in of TFA:TIPS:EDDT:H_2_O 90:2.5:2.5:5 (trifluoroacetic acid : triisopropyl silane : 2,2’-(Ethylenedioxy)diethanethiol: water) and precipitated in 10 volumes of ice cold Et_2_O, collected by centrifugation, and washed with the same volume of ice cold Et_2_O, before purification by preparative HPLC. The peptides were then lyophilized, resuspended in ultrapure water, neutralized (to pH 6.0 to avoid cysteine oxidation) with dilute NaOH, sterilized by filtration at 0.2 μm, aliquoted and re-lyophilized. Dry aliquots were stored at −20 °C or below.

### Synthesis of DCMAC-SAG-SH

0.2 mmol of SAG-SH with the sequence **GGGGLERCL**-NH2, still bound to rink-amide resin and bearing standard side chain protecting groups (OtBu on glutamate, Pbf on Arginine, Trt on cysteine) and with its N-terminal Fmoc removed, was caged by introducing 0.8 mmol DCMAC-NPC (1d) dissolved in 4 ml of DMF:Pyridine 10:1 in the reaction vessel. The reaction was left to proceed overnight in the dark with shaking. The substitution was found to be quantitative at this point with analytical HPLC monitoring on microcleavages. After extensive washing of the resin-supported caged peptide with DMF and DCM and nitrogen-flow drying, the peptide was cleaved, deprotected, washed, purified, neutralized, sterilized and stored as described above. The identity of the caged peptide was confirmed by MALDI-TOF. [M+H]^+^ expected/found: 1195.

### Synthesis of DEAC derivatives (2a, 2b, 2c, 2d, DEAC-SAG-SH)

7-diethylamino 4-methylcoumarin (DEAC, 2a) was purchased from TCI Deutschland GmbH.

The following derivatizations followed the same procedures as described for DCMAC derivatives, with comparable yields.

### Synthesis of 6-bromo 7-hydroxy 4-chloromethyl coumarin (BHC-Cl, 4a)

2 g (10.6 mmol) of 4-bromoresorcinol were dissolved in 25 ml of methanesulfonic acid. 2.2 mL of ethyl 4-chloroacetoacetate (16 mmol) was added dropwise and the reaction was stirred for 2 h at room temperature. The mixture was then precipitated by dropwise addition into 500 mL of ice-cold water with vigorous stirring, and left standing on ice for 30 min. The precipitate was recovered by vacuum filtration, washed 5 times with 50 mL of cold water, and dried overnight at 50 °C under high vacuum, to yield 2.9 g of the product (94% yield), as an off-white solid.

### Synthesis of 6-bromo 7-hydroxy 4-hydroxymethyl coumarin (BHC-OH, 4b)

1 g (3.5 mmol) of BHC-Cl (4a) was dissolved in 30 ml of DMF, and 15 ml of 1 M HCl were added. The mixture was refluxed at 95 °C under nitrogen for 24 h, and concentrated on a rotary evaporator at 65 °C under reduced pressure. The residue was resuspended in ≈20 ml of DMF and purified by preparative HPLC to yield 0.63 g of the product as a white solid (67% yield).

### Synthesis of (6-bromo 7-hydroxy coumarin 4-yl)methyl (4-nitrophenyl) carbonate (BHC-NPC, 4c)

63 mg (0.23 mmol) of BHC-OH (4b) and 55 mg (0.28 mmol) of NPC were reacted in 500 μl of pyridine on ice with vigorous stirring. The reaction was monitored by analytical HPLC. The resulting solution of BHC-NPC was used directly in the following step.

### Synthesis of BHC-SAG-SH

Freshly activated BHC-NPC (4c), ≈0.2 mmol in pyridine solution, was reacted with 0.05 mmol of solid-phase supported SAG-SH (after N-terminal Fmoc removal, side-chains still protected). The reaction was left to proceed overnight at room temperature with vigorous shaking. The peptide was then cleaved and purified as described above.

### Synthesis of NPPOC-SAG-SH

0.05 mmol of solid-phase supported SAG-SH (after N-terminal Fmoc removal) was reacted with 50 μl (0.13 mmol) of 2-(2-Nitrophenyl)propyl chloroformate (NPPOC-Cl, 3) in the presence of 34 μl (0.2 mmol) ^i^Pr_2_NEt in 4 ml of DMF. After 15 minutes, the reaction was found to be complete and the cleavage and purification were done as described above.

### Synthesis of DCMAC-SAG-NHS

A DCMAC-SAG peptide of sequence DCMAC-**GGGK**-NH2 was cleaved from Rink-amide resin and deprotected as described above. The peptide identity was confirmed by LC-MS: [M+H]^+^ expected = 649.46 found = 649.6. Then, 100 mg (0.15 mmol) of peptide in 5 ml of dry DMSO were added dropwise to 200 mg (0.78 mmol) of disuccinimidyl carbonate (DSC) in 4 ml of dry DMSO. Monitoring on analytical HPLC monitoring showed around 10-15% peptide substitution. After dropwise addition of 27 μl (0.15 mmol) of ^i^Pr_2_NEt, monitoring showed around 80% peptide substitution, and an additional 27 μl of ^i^Pr_2_NEt showed complete conversion. The product was purified by preparative HPLC on a gradient from 5 to 50% ACN in water+0.1% TFA in 25 min, to yield 82.5 mg (68%) of DCMAC-**GGGK**(N-hydroxysuccinimide carbamate)-NH2 (DCMAC-SAG-NHS), as confirmed by LC-MS: [M+H+] expected/found = 790.7.

The peptide was then lyophilized, resuspended in water, sterilized by filtration, re-lyophilized in small aliquots (1.6 mg, 0.2 μmol), and stored at −20 °C.

### Synthesis of DCMAC-SAG-TGQ

20 mg (15 μmol) of TG glutamine donor peptide (TGQ) with sequence **NQEQVSPLERCG**-NH_2_ were reacted with 4 μl (40 μmol) of divinyl sulfone in 1 ml of TEOA buffer (triethanolamine 300 mM, pH 8.0). After 30 s, test with Ellman’s reagent showed all the thiols were reacted and the reaction was stopped by freezing in liquid nitrogen. The vinyl sulfone-peptide conjugate was purified by preparative HPLC. The collected fractions were lyophilized, and resuspended in TEOA buffer for further reaction with 2.5 mg of DCMAC-SAG-SH (2 μmol). The reaction appeared to be complete after 15 min with analytical HPLC monitoring, and the conjugate (as well as unreacted excess TGQ for recycling) was HPLC-purified, lyophilized, resuspended in water, neutralized, sterilized by filtration, aliquoted, re-lyophilized, and stored at −20 °C.

Conjugation of the DCMAC-SAG-TGQ peptides at 400 μM to 0.83% hyaluronan bearing the other TG ligand (TGK, sequence Ac-**FKGGERCG**-NH_2_) in the presence of 20 U/ml TG (Fibrogammin, CSL Behring) was studied by GPC, and the conjugation was found to happen swiftly, with a time constant of 2.4 to 2.6 min but until a limit of approximately 50% coupling. Further attempts to form a gel from these pre-reacted DCMAC-SAG-TGQ and hyaluronan did not work, showing that the limit is due to quick loss of enzymatic activity of TG in the presence of its substrate peptides. As a result, DCMAC-SAG-TGQ peptides were not pre-reacted but rather added to the other gel precursors and coupled to the polymer backbone simultaneously with gelation.

### Synthesis of DCMAC-TGK-SH

The peptide Ac-**FK**(DCMAC)**GGERCG**-NH2 (DCMAC-TGK-SH) was synthesized as a proof-of-concept of caging on arbitrary amino groups other than the N-terminus of the peptide. We synthesized with standard methods the peptide Ac-**FKGGERCG**-NH2 using a lysine with a hydrazine-sensitive orthogonally-protected side chain, Fmoc-Lys(Ddiv)-OH. The other protecting groups were standard acid-labile groups, i.e. fmoc-Glu(tBu)-OH, Fmoc-Cys(Trt)-OH, Fmoc-Arg(Tbf)-OH. The resin-supported protected-peptide, Ac-**FK**(Ddiv)**GGE**(tBu)**R**(Tbf)**C**(Trt)**G**-(Rink Amide), was then treated with 2% hydrazine in DMF until complete Ddiv removal, exposing the side chain amino group of the lysine residue. DCMAC-NPC coupling and peptide cleavage and deprotection were then performed as described above, and the identity of DCMAC-TGK-SH was confirmed by MALDI-TOF: [M+H+] expected/found - 1185. Of note, when acetylation of the N-terminus is unwanted, terminal fmoc should still be removed and replaced by a TFA-labile tert-butyloxycarbonyl (BOC) protecting group before Ddiv deprotection and cage conjugation, as carbamate linked two-photon active cages do not withstand piperidine:DMF treatment used for fmoc removal.

### Synthesis of alginate-vinyl sulfone

1.17 g (6 mmol) of MES were dissolved in 40 ml of water, followed by 400 mg (2 mmol of the repeat unit) of high molecular weight sodium alginate (Sigma A7128), dissolved overnight with gentle rocking. The pH of the resulting solution was 4.5. We then added 11.9 mg (0.05 mmol) of of 3,3’-dithiobis(propanoic dihydrazide) (DTPHY, Frontier scientific), and 19.2 mg of (0.1 mmol) of EDC (Fluka) predissolved in 1 ml of water and added dropwise over vigorous stirring. Stirring was then stopped, and the reaction was left to proceed overnight. The disulfides were then reduced with 57.3 mg (0.2 mmol) of tricarboxyethyl phosphine hydrochloride (TCEP, Fluorochem), predissolved in 1 ml of water and added dropwise while swirling the flask. The reduction was left to proceed overnight in the sealed standing flask at RT. The resulting thiolated alginate solution was supplemented with 2 g of NaCl, and dialyzed against ultrapure water balanced to pH 4.5 by addition of HCl. After 24 h, the solution was recovered and added dropwise over stirring to 0.5 ml (10 mmol) of divinyl sulfone in 10 ml of TEOA. Special care was taken as divinyl sulfone is fatal in contact with skin. After 3 h of reaction at room temperature, the solution was supplemented with 4 g NaCl and dialyzed against ultrapure water. After completion, the alginate-vinyl sulfone sodium salt solution was sterilized by filtration, aliquoted, lyophilized, and stored at −20 °C. The structure of the vinyl sulfonated polymer was confirmed by NMR, and the substitution ratio was measured with a modified version of the Ellman’s assay. A 10 mg/ml alginate resuspended stock in TEOA buffer was reacted in equal proportions with a 3 mM β-mercaptoethanol solution in the same buffer. A control without alginate-vinyl sulfone was included in parallel. The Michael addition was left to proceed for 5 h at room temperature, and the remaining thiols were quantified with a classical Ellman’s assay. We found 1.6 mM of thiols in this 10 mg/ml stock, which corresponds to 3.2% substitution.

### Synthesis of TG cross-linkable hyaluronan

The synthesis was done as described previously^39,50^.

### Synthesis of 2-((2-(vinylsulfonyl)ethyl)thio)acetic acid (VS-COOH, 5a)

50 μl (0.7 mmol) of thioglycolic acid, 140 μl (1.4 mmol) of divinyl sulfone, and 7 μl (0.05 mmol) of DBU were reacted in 1 ml DMF for 30 min on ice. The reaction mixture was then diluted to 10 ml with 30 mM aqueous HCl, and purified by HPLC (5% ACN for 10 min, 5 to 90 over 20 min), followed by lyophilization, to get the product in quantitative yield (114 mg) as a white crystalline solid.

### Synthesis of 2,5-dioxopyrrolidin-1-yl 2-((2-(vinylsulfonyl)ethyl)thio)acetate (VS-NHS, 5b)

160 mg (0.76 mmol) of VS-COOH were dissolved in 8 ml of DCM, and reacted with 180 μl (1.14 mmol) of DIC and 132 mg (1.14 mmol) of NHS for 10 min at RT. Monitoring on thin layer chromatography (TLC) in EtOAc with KMnO_4_ staining showed the reaction was complete (Rf = 0.64). The product was purified by flash chromatography in EtOAc, and the solvent was removed under reduced pressure, to get 191 mg (82% yield) of product as a white crystalline solid.

### Synthesis of DCMAC-SAG-fibrinogen

40 mg of bovine fibrinogen powder (Sigma F86030), which include 30 mg (88.2 μmol) of actual protein, were dissolved in 1 ml of PBS (Gibco, calcium and magnesium free, pH7.4). 154 μg (500 nmol) of VS-NHS were then taken as 3.08 μl of a 5% stock in DMSO, prediluted in 100 μl of PBS, and the fibrinogen solution was added to the VS-NHS solution and mixed swiftly. The mixture was let to react for 20 min before adding 770 μg (650 nmol) of DCMAC-SAG-SH, pre-dissolved in 200 μl of TEOA buffer. As peptides typically contain some salts, the exact peptide molarity was measured by Ellman’s assay rather than only on a microbalance. The reaction was left to proceed for 2h at 37°C, sterilized by filtration, and the modified fibrinogen was then precipitated by adding 2 volumes of sterile brine, and recovered by centrifugation at 2000 g for 2 min. After further washing with brine:water 1:1, the pellet was resuspended in 1.5 ml of sterile PBS (keeping the suspension at 37 °C until it clarifies), and the protein and cage concentrations were measured on a spectrophotometer, using calibration curves for the absorbance at 280 nm of fibrinogen solutions, and for the fluorescence at 400/470 nm of DCMAC-SAG-SH. The final stocks were found to have 15 mg/ml of fibrinogen (75% recovery) with 187 μM of DCMAC-SAG substitution (4.2 caged peptides/protein, 75% efficiency in the substitution reaction).

### Synthesis of Avidin-SAT-F

300 μg (4.41 nmol) of Avidin (Thermo) stored frozen in 150 μl of water were melted and reacted with 16.2 μg (53 nmol) of VS-NHS (the 5% VS-NHS stock in DMSO was prediluted 100-fold in PBS, and 32.4 μl of this solution were added) and 21 μg (45 nmol) of fluorescein-NHS (Thermo, from a 1% fluorescein-NHS stock in DMSO stored frozen, prediluted 50 fold in PBS before addition of 105 μl). The reaction was left to proceed for 30 min at RT, before adding 650 μg (462 nmol) of a SA threonine-containing substrate peptide (SAT) with the sequence Ac-**GCREDDDLPMTGG**-NH_2_ pre-dissolved in 100 μl of TEOA buffer (300 mM, pH 8.0). The reaction was left to proceed further for 1 h at 37 °C, and the Avidin-SAT-F was purified by GPC using Tris 50 mM + NaCl 300 mM balanced to pH 7.5 as the running phase. The collected fractions were diluted two-fold with ultrapure water to balance the osmolarity, sterilized by filtration, and re-concentrated with vivaspin protein concentrators (GE Healthcare, 10 000 MWCO) to recover 500 μl of stock at 0.4 μg/μl of protein content (approx. 60% protein recovery), with 6 SAT peptides/avidin tetramer (as quantified on MALDI-TOF, Fig. S5). This stock was stored frozen in small aliquots until use.

### Synthesis of NGF-PEG-Biotin

80 μg of NGF in water solution at 1 μg/μl was reacted with 1.2 equivalents (over the dimer) of NHS-PEG_12_-Biotin (Thermo), pre-dissolved in 32 μl of PBS. The reaction appeared complete after 2 h according to HPLC monitoring, and the stock was aliquoted and frozen without further purification. The substitution ratio (60% of the monomers, 1.2 Biotin per dimer) was measured on HPLC and confirmed by MALDI-TOF mass spectroscopy.

### BSA-rhodamine

10 mg of bovine serum albumin (BSA) in 500 μl of carbonate buffer, 100 mM, pH 9.5 were reacted with 20 μl of a DMSO stock of 10 mg/ml rhodamine-isothiocyanate (RITC). The protein was purified by GPC and re-concentrated with a Vivaspin protein concentrator, 10 000 MWCO. The 5 mg/ml resulting stock was stored at −20°C.

### Alginate gels with caged peptides

Vinyl sulfonated alginate and unmodified alginate, 1% stocks in TEOA buffer (300 mM, pH 8.0), were combined in order to have a 1% alginate stock with 500 μM vinyl sulfone handles. This stock was further reacted by Michael addition with NPPOC-SAG-SH, BHC-SAG-SH, DEAC-SAG-SH, and DCMAC-SAG-SH, with the peptides added at 1 mM from dry powder and reacted overnight.

To form gels for bulk non-specific adsorption measurements, 10 μl drops of these 1% functionalized alginate stocks (or non-functionalized control alginate, or mixtures of these to achieve lower molarities of functional groups) were deposited on top of 100 μl of CaCl_2_ solution in each well of a 96 well plate. The gels were further washed twice one hour with the CaCl_2_ solution in order to remove unbound peptide.

For two-photon patterning experiments, 1% alginate stocks with 100 μM of caged peptides were cast into polydimethylsiloxane (PDMS) cylindrical molds of 500 μm height and 3 mm inner diameter adhered to glassbottom microscopy chambers, and precoated with poly-L-lysine (PLL, 1 mg/ml inserted in the PDMS mold for 10 min followed by washing with ultrapure water and drying). A cellulose filter membrane pre-wetted with 100 mM CaCl_2_ was then deposited on the top surface, and an additional 100 μl of 100 mM CaCl_2_ added on top. Gelation was left to proceed for 1 h, before covering the gels with a 1:1 mixture of 100 mM CaCl_2_ and Tris buffered saline (TBS, Tris 50 mM, NaCl 150 mM, pH7.5) until photopatterning.

### BSA adsorption study

Functionalized alginate gels were incubated for 1 h in 1 mg/ml BSA-rhodamine solution (mixing the protein stock and CaCl_2_ solutions 1:4). They were then washed with a 1: 1 mixture of TBS and CaCl_2_ 100 mM, for 48 h in total with frequent buffer changes. Finally, the fluorescence of the gels was measured on a plate reader (excitation 555 nm emission 590 nm). A calibration curve from serial dilutions of BSA-rhodamine in the same buffer was also constructed in order to convert the fluorescence values to adsorbed protein concentrations. Control gels with the various caged peptides or no functionalization were also measured, without incubation with the fluorescent rhodamine, and found to have no fluorescence at these wavelengths that would compromise the assay.

### Photo-patternable collagen gels

An aliquot of DCMAC-SAG-NHS (1.6 mg, 0.2 μmol) was resuspended in 20 μl of dry DMSO to form a 1 mM stock. Patternable collagen was formed by mixing 50 μl of ice-cold collagen acidic stock (AteloCell, bovine dermis collagen, 5 mg/ml), 1 μl of DCMAC-SAG-NHS DMSO stock, and 20 μl of 5% w/v NaHCO_3_ for neutralization and balancing of osmotic/ionic strength. The mixture was mixed vigorously while still standing on ice, and cast into PDMS molds. Gelation was left to proceed for 10-20 min at 37 °C, 5% CO_2_. The gels were then covered with TBS to wash unbound peptides and avoid drying during the following photopatterning process. Fluorescence measurements on a plate reader against a calibration curve from serial dilutions of DCMAC-SAG were used to determine the substitution of collagen gels, and the final bound peptide concentration was found to be of ≈ 145 μM.

### Photo-patternable basement membrane extract (Matrigel)

An aliquot of DCMAC-SAG-NHS (1.6 mg, 0.2 μmol) was resuspended in 1 ml of sterile water, neutralized with dilute NaOH using a small amount of phenol red as a pH indicator, and sub-aliquoted in portions of 0.16 mg (0.02 μmol), snap-frozen in liquid nitrogen, and re-lyophilized for storage at −20 °C. 0.02 μmol aliquots were then individually resuspended in 80 μl of ice-cold Matrigel (Corning, 9 mg/ml in Dulbecco’s modified Eagle’s medium), and cast in PDMS molds. After 15 min of gelation at 37 °C, the gels were washed with PBS (incl. calcium), and used for two-photon patterning as described below. Quantification of DCMAC-SAG coupling was performed using full gel fluorescence readings on a plate reader (gels washed overnight were loaded between the windows of a 1 mm pathlength Take3 plate) against a calibration curve from serial dilutions of peptide as described above, and we found that the gels were substituted with 200 μM of caged peptides.

### Photo-patternable Hyaluronan gels

The gelling mixture consisted of DCMAC-SAG-TGQ at 200 μM, TG cross-linkable hyaluronan at 0.75% w/v and activated factor XIII (TG, Fibrogammin from CSL Behring) at 20 U/ml, in a buffer consisting of 100 mM glucose, 50 mM Tris, 50 mM CaCl_2_, pH 7.5. Gelation followed within 1 minute, and the gels were then covered with TBS after 10-15 min and until patterning.

### Photo-patternable fibrin gels

DCMAC-SAG-fibrinogen and thrombin (Tisseel, Baxter) stocks were combined in TBS + 1 mM CaCl_2_ for final concentrations of fibrin 3 mg/ml and thrombin 0.5 U/ml. Gelation was left to proceed for 15 min at 37 °C, and the gels were then covered with TBS.

### DRG isolation and embedding (for two-step patterning, Fig. 6a-e)

Dorsal root ganglia (DRGs) were isolated from E9-10 chick embryo, and collected in PBS on ice. Immediately before preparing the gels, they were washed in a 0.75% solution functionalized TG-cross-linkable hyaluronan, to avoid the formation of a thin buffer layer between the DRGs and their matrix. Number 5 forceps (FST) were used to transport DRGs, coated with albumin (by dipping in a 5% solution in PBS) to avoid DRG attachment. Gels were casted as described above, and one DRG placed in the center of each gel before gelation occurs.

The DRGs were then cultured at 37 °C, 5%CO_2_, in neurobasal medium with B27, glutamax and penicilin/streptomycin. Medium was added immediately after gel formation and when applicable two-photon patterning, and changed twice during the first 24 h to remove uncoupled protein. Half of the medium was then renewed once a week.

### Two-photon patterning (2PP, Fig. 3a, 4b, 5e-h)

The glass-bottom chambers with gels formed with one of the five materials listed above were loaded on a Leica SP8 multiphoton microscope equipped with a spectra physics Mai Tai laser, 25x water immersion objective and LASX software, or a Leica SP5 multiphoton microscope equipped with a Coherent Chameleon laser, LAS-AF software and 20x water immersion objective. The laser intensity and scan number were varied as indicated in each figure legend. The step in the z direction was kept constant at 1 μm, and the pixel size in the xy plane at 0.5 μm. These dimensions approximately fit the size of the point spread function and ensure the formation of continuous patterns. The scanning speed was kept constant at 600 Hz.

In order to pattern complex shapes or shades, we programmed custom Matlab scripts which take standard .tif pixel data or .stl 3D mesh data, and generate instructions matching the format of the Leica live data mode instructions. The 3D rose model used as a test file was created in Blender, exported as a binary .stl file, and reexported as an ASCII .stl file using Meshlab. The scripts use a live data mode file exported from the microscope as a template for the formatting and an input for the microscope parameters (detector gains, lasers, filter positions etc). 3D structures (from .stl) are made layer by layer, using the region of interest (ROI) scanning possibility of the microscopes, which physically consists in fast light switching on and off by an electro-optical modulator (EOM). For shades of greyscale patterns, the EOM/ROIs are used as well, and we sequentially scan a series of ROIs (grey level masks) at increasing laser intensity. Calibration curves of pattern intensity vs uncaging power were taken into account in order to produce a linear relation between the reference grey level and the final patterned avidin concentration.

Immediately after photopatterning, the buffer covering the gels was removed and the gels were covered with an incubation solution consisting of 0.7 μg/μl avidin-SAT-F stock (80%), biotinylated NGF stock (5%), SA 80 μM stock (5%) and 100 mM CaCl_2_ stock (10%). The coupling to the uncaged areas was left to proceed for 30-40 min, and the gels were then transferred to cell culture medium (with cells) or TBS.

### One-step 2PP (Fig S4c, 5a-d, 6f-k)

One-step patterning was demonstrated in the hyaluronan and fibrin gels. The process is the same as for the two-step patterning, except that all the components normally added in the incubation solution are directly mixed with the gel precursors. The stocks combined for hyaluronan gels are: TG cross-linkable hyaluronan 3%, DCMAC-SAG-TGQ 4 mM, Avidin-SAT-F 0.4 μg/μl + NGF-biotin 35 ng/μl, SA 80 μM, and activated TG 200 U/ml. The percentage of these stocks in the mix are respectively: 25:10:50:5:10. The TG was added last to trigger the gelation. Freshly isolated DRGs were encapsulated within 30 s after gel casting when applicable. The stocks combined for fibrin gels were: DCMAC-SAG-fibrinogen 10 mg/ml, SA 80 μM, avidin-SAT-F 0.4 μg/μl + NGF-biotin 35 ng/μl, thrombin 10 U/ml, and CaCl_2_ 100 mM, in proportions of 30:5:50:5:10. Thrombin was added last to trigger the gelation. Gelation occurred within minutes, and the gels were then immediately photo-patterned as described above, and then transferred to TBS (without cells) or cell culture medium (with cells).

### Immunocytochemistry

Hydrogels were fixed for 1 h in ice-cold 4% formaldehyde solution in PBS, washed with PBS for at least 1 h, blocked with 5% bovine serum albumin (BSA) in PBS overnight, incubated with primary antibody for (Tuj1, Sigma T5076, 1:500 in PBS+3% BSA), washed with PBS 2 x 1h and 1x overnight, incubated with secondary antibody (goat anti-mouse Alexa 594, Thermo, 1:400 in PBS+3%BSA), washed with PBS 2×1h and 1x overnight, and stained with DAPI and phalloidin-tetramethylrhodamine (thermo) for 2 h, then washed with PBS and transferred on a coverslip for imaging with confocal/two-photon Leica SP8 microscopes with 20x or 25x water objectives.

## Supplementary figures

**Figure S1.**
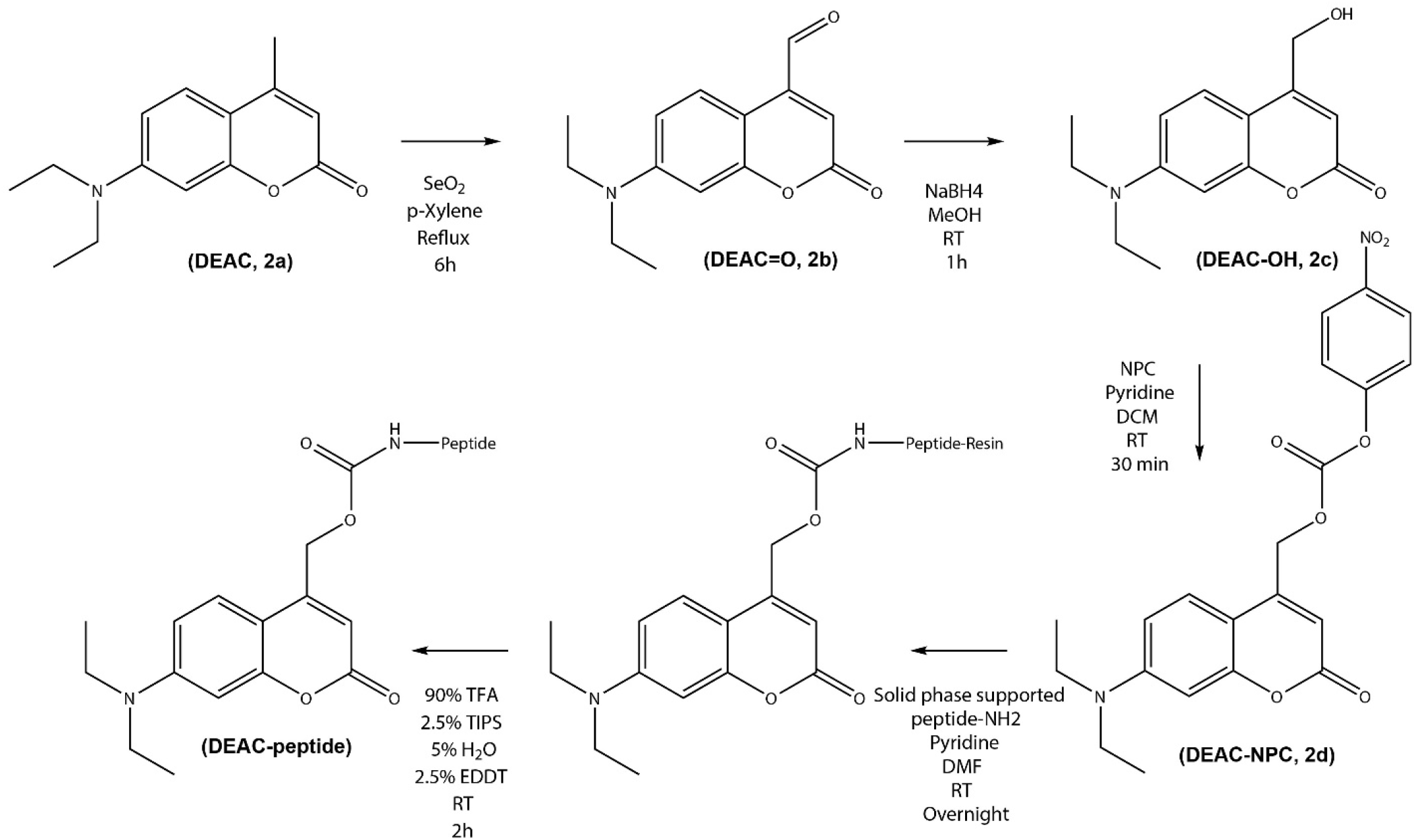
Synthesis of 7-diethylaminocoumarin (DEAC) caged peptides.

**Figure S2.**
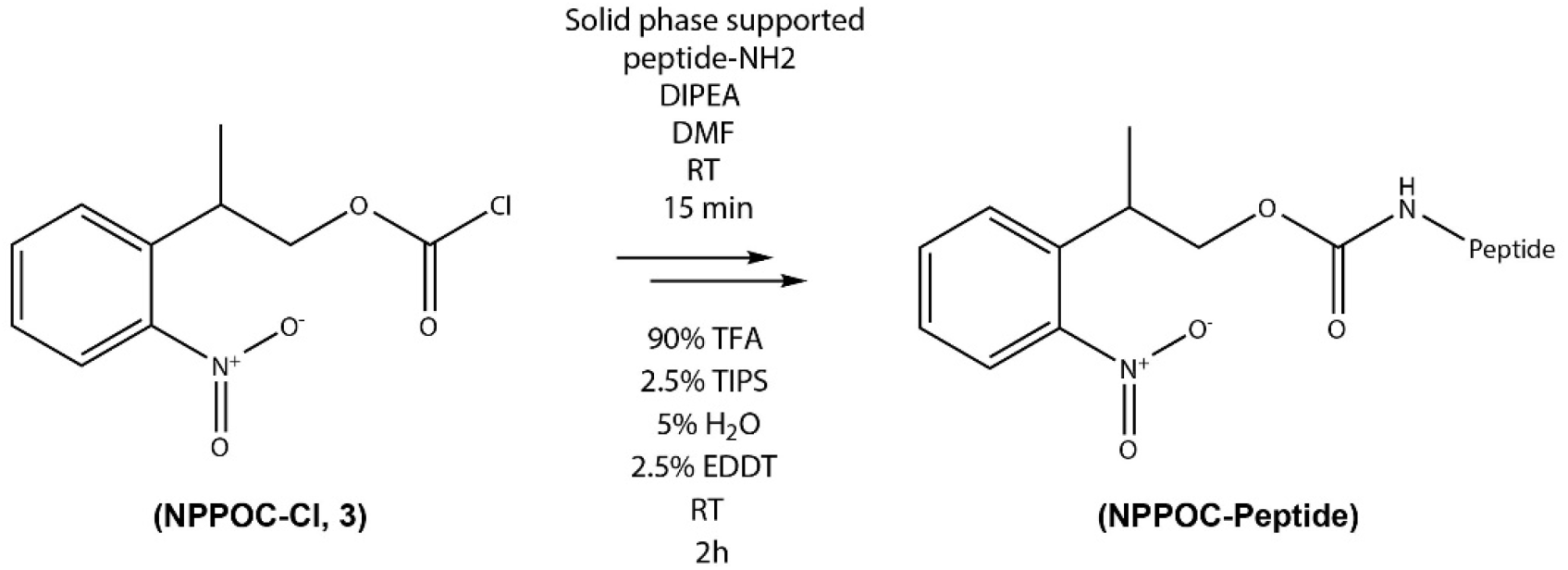
Synthesis of 2-(2-Nitrophenyl)propyl carbamate (NPPOC) caged peptides.

**Figure S3.**
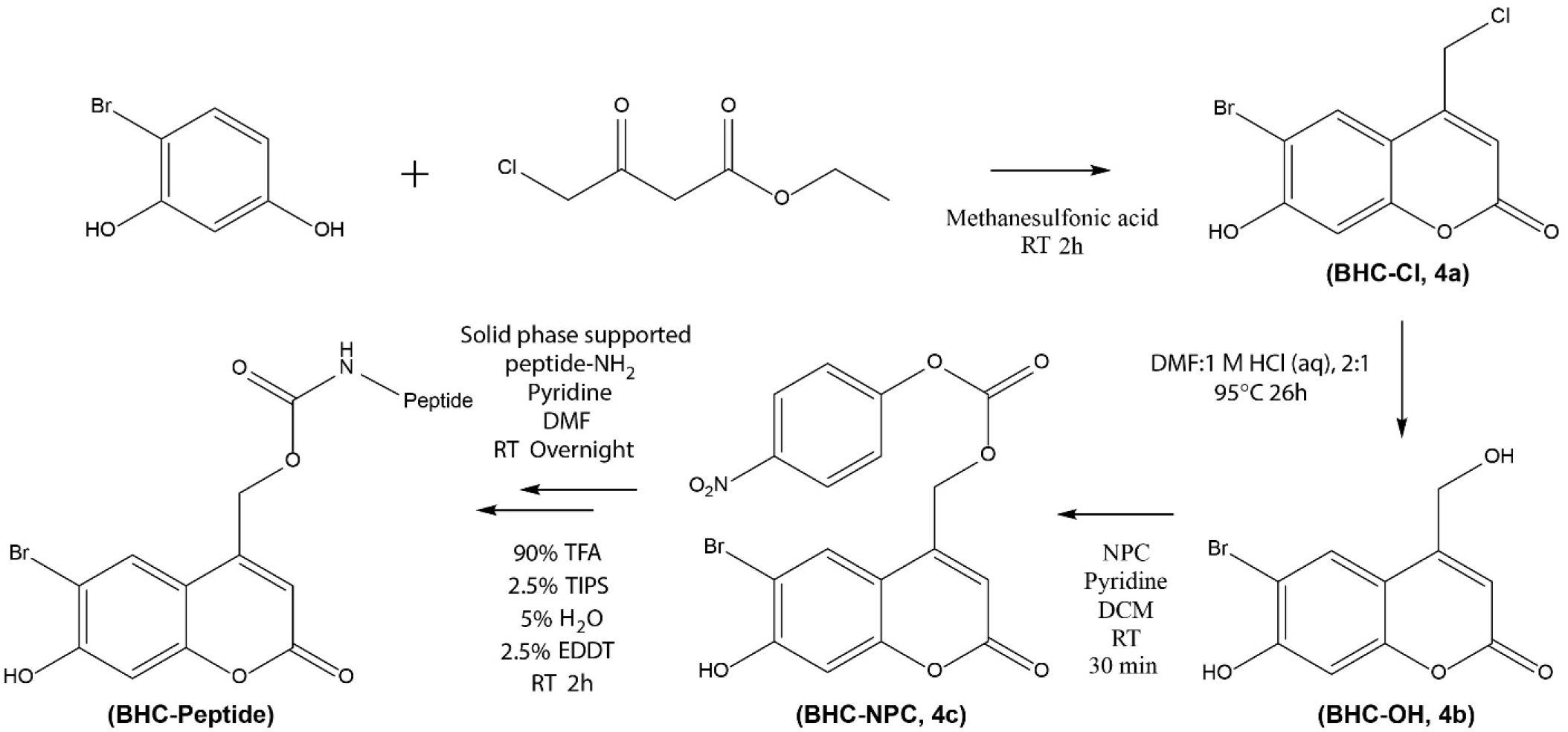
Synthesis of 6-bromo 7-hydroxycoumarin (BHC) caged peptides.

**Figure S4.**
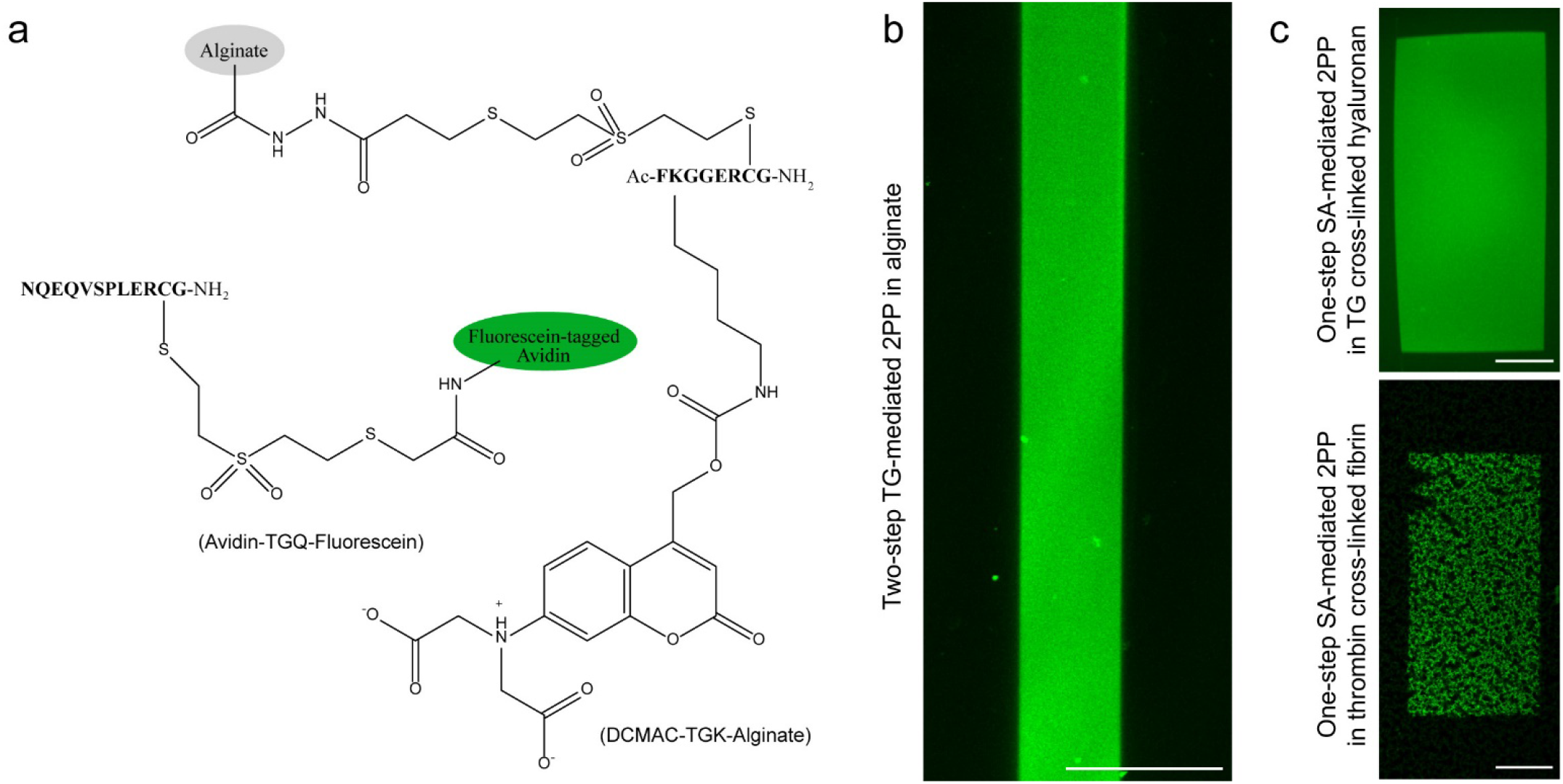
Two-step human activated factor XIII (TG) mediated 2PP of avidin in alginate, using DCMAC-caged substrate peptides, and one-step SA-mediated 2PP in biological matrices, utilizing the orthogonality of SA to TG and thrombin. (**a**) Description of the chemicals used for TG-mediated 2PP, including alginate functionalized with caged TG-Lysine donor peptides (TGK) and avidin functionalized with TG-Glutamine donor peptides (TGQ). (**b**) Example of TG-mediated 2PP outcome. (**c**) One-step SA-mediated 2PP: resulting patterning in hyaluronan and fibrin gels, using 30% laser power and 1-2 scans respectively, on a Leica SP8 inverted microscope with 25x water immersion objective. Scale bars: 100 μm.

**Figure S5.**
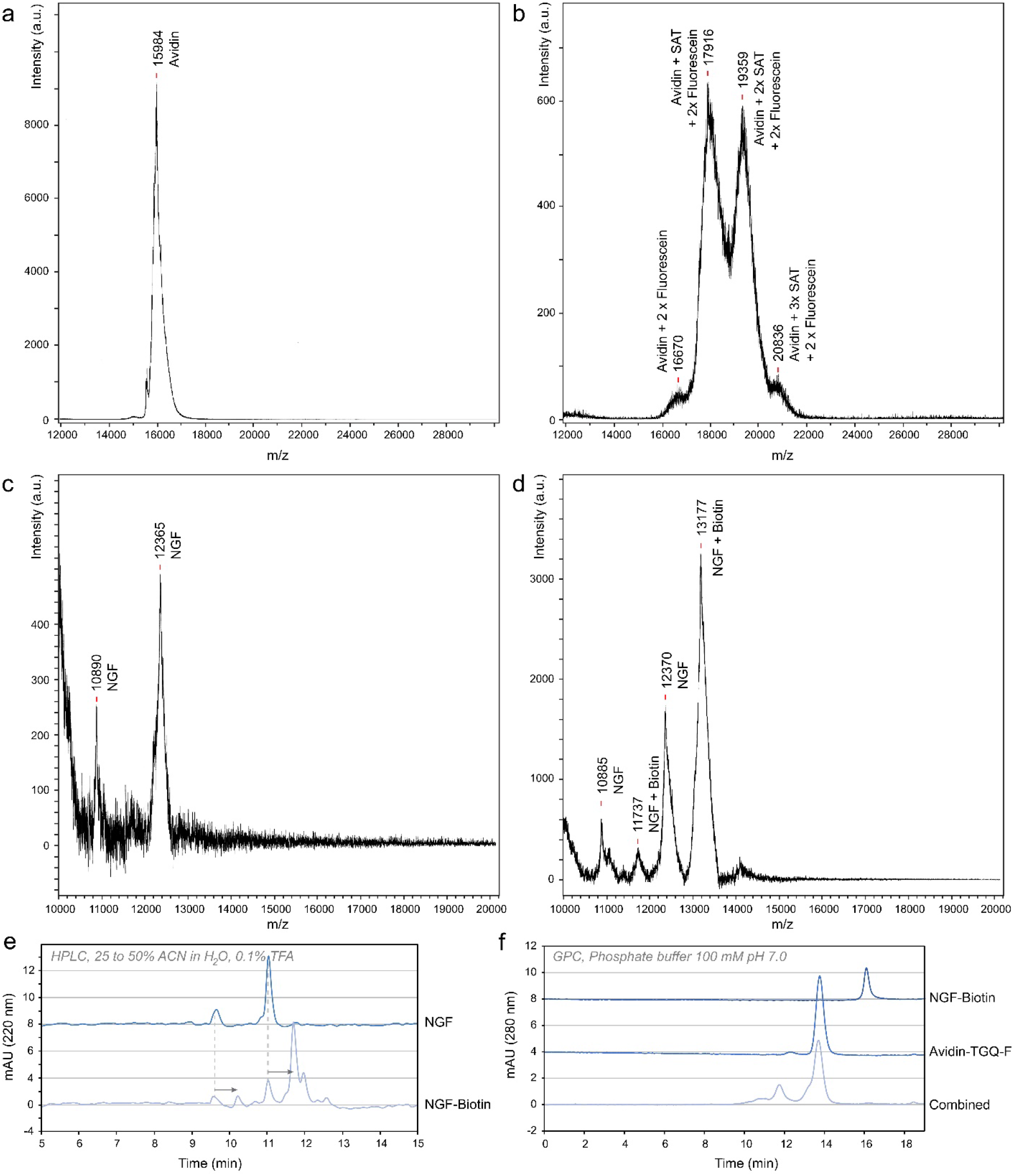
Characterization of SAT and fluorescein-substituted avidin (Avidin-SAT-F) and biotinylated NGF (NGF-Biotin). (**a, b**) Matrix-assisted laser desorption and ionization time-of-flight mass spectroscopy (MALDI-TOF) of avidin and SAT-substituted avidin, showing approximately 1.5 substitutions on average per monomer, i.e. 6 substitutions per avidin tetramer. (**c,d**) MALDI-TOF of NGF-Biotin, showing approximately two thirds of the NGF monomer bear a biotin (with PEG spacer), i.e. ≈1.3 per NGF dimer. (**e**) Confirmation of the substitution of NGF with biotin on C18 analytical HPLC. (**f**) Confirmation of NGF-Biotin binding to modified avidin by gel permeation chromatography (GPC).

**Figure S6.**
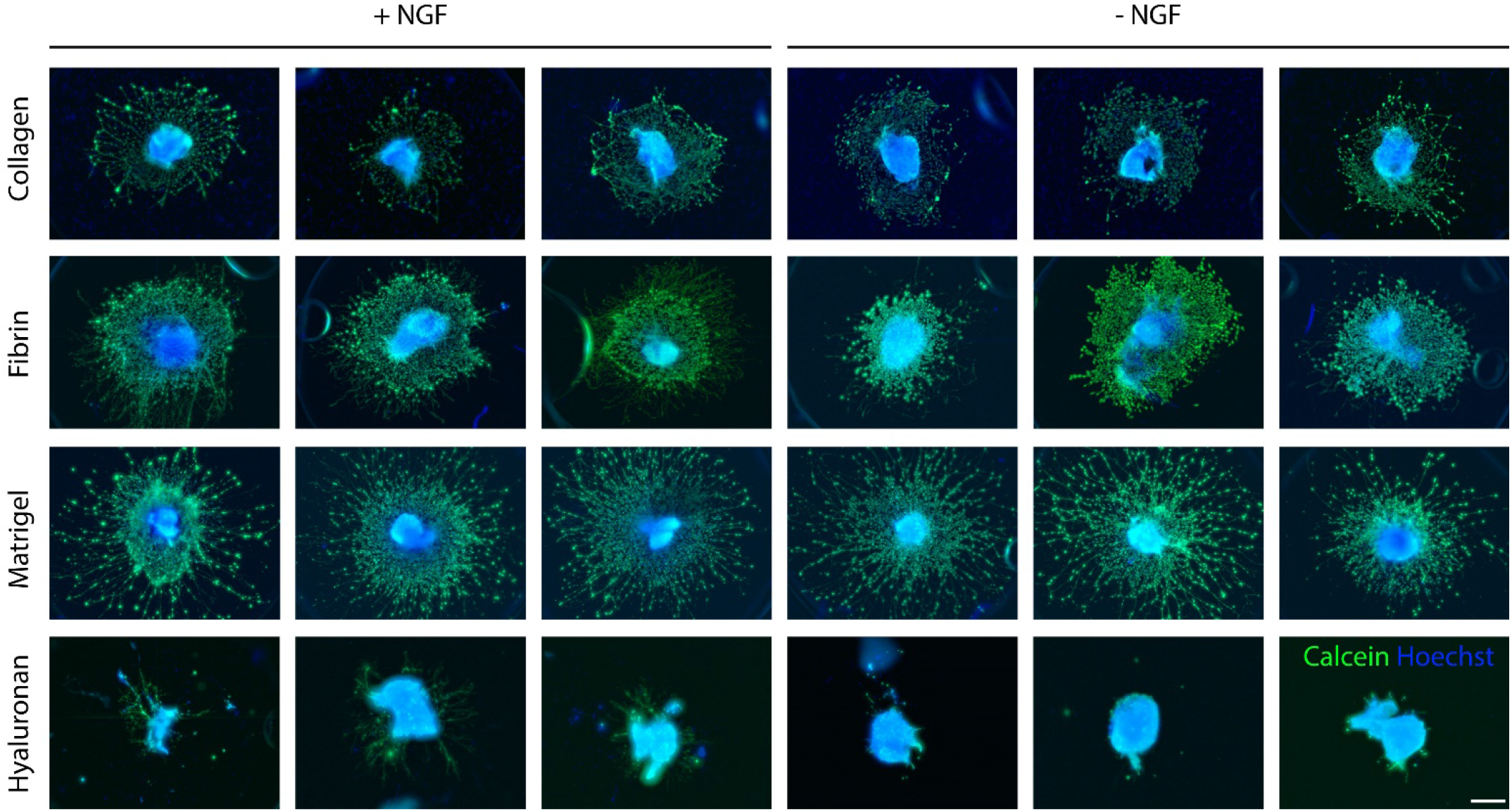
Comparison of invasion by chick E9 DRGs axons and glia after 4 days in culture in various biological matrices, in the presence or absence of NGF at 200 ng/ml. Maximum intensity projection of widefield fluorescence images of calcein-AM (live cells and processes) and Hoechst (nuclei), with background subtracted and gamma adjusted to 0.3 to facilitate axon visualization. Scale bar: 500 μm.

**Figure S7.**
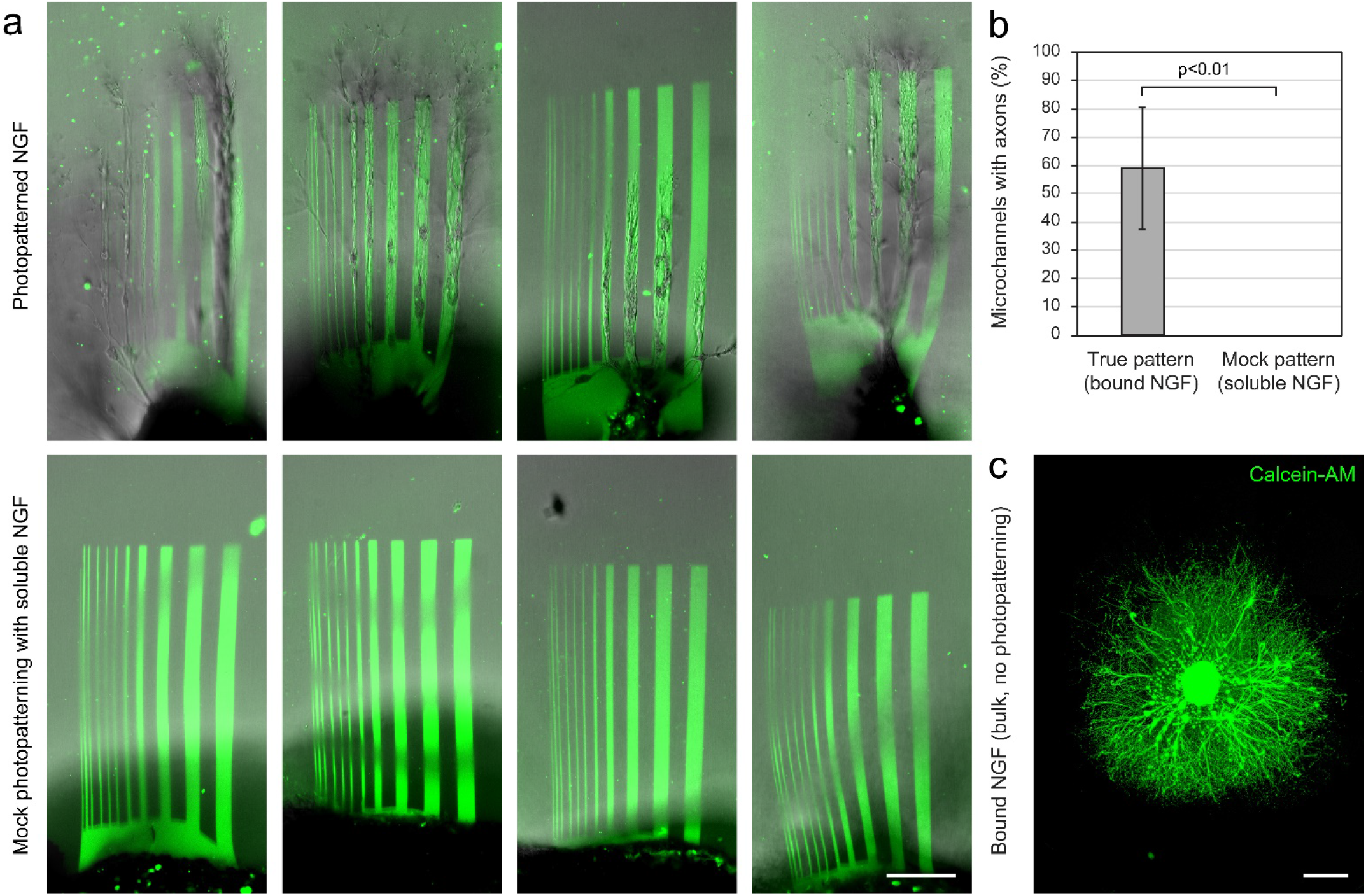
The axonal growth and guidance in hyaluronan hydrogels are specifically mediated by immobilized NGF. (**a-b**) Comparison of the axon extension in standard 2PP experiments vs experiments replacing NGF-Biotin by soluble (non-biotinylated) NGF. Each image is from a replication on a different DRG, after 2 days in culture. Scale bar: 100 μm (**c**) Axonal growth from a chick E9 DRG in a hydrogel with NGF anchored in bulk (by forming the hydrogel in the presence of avidin-TGQ and NGF-Biotin). Scale bar: 500 μm.

## NMR spectra

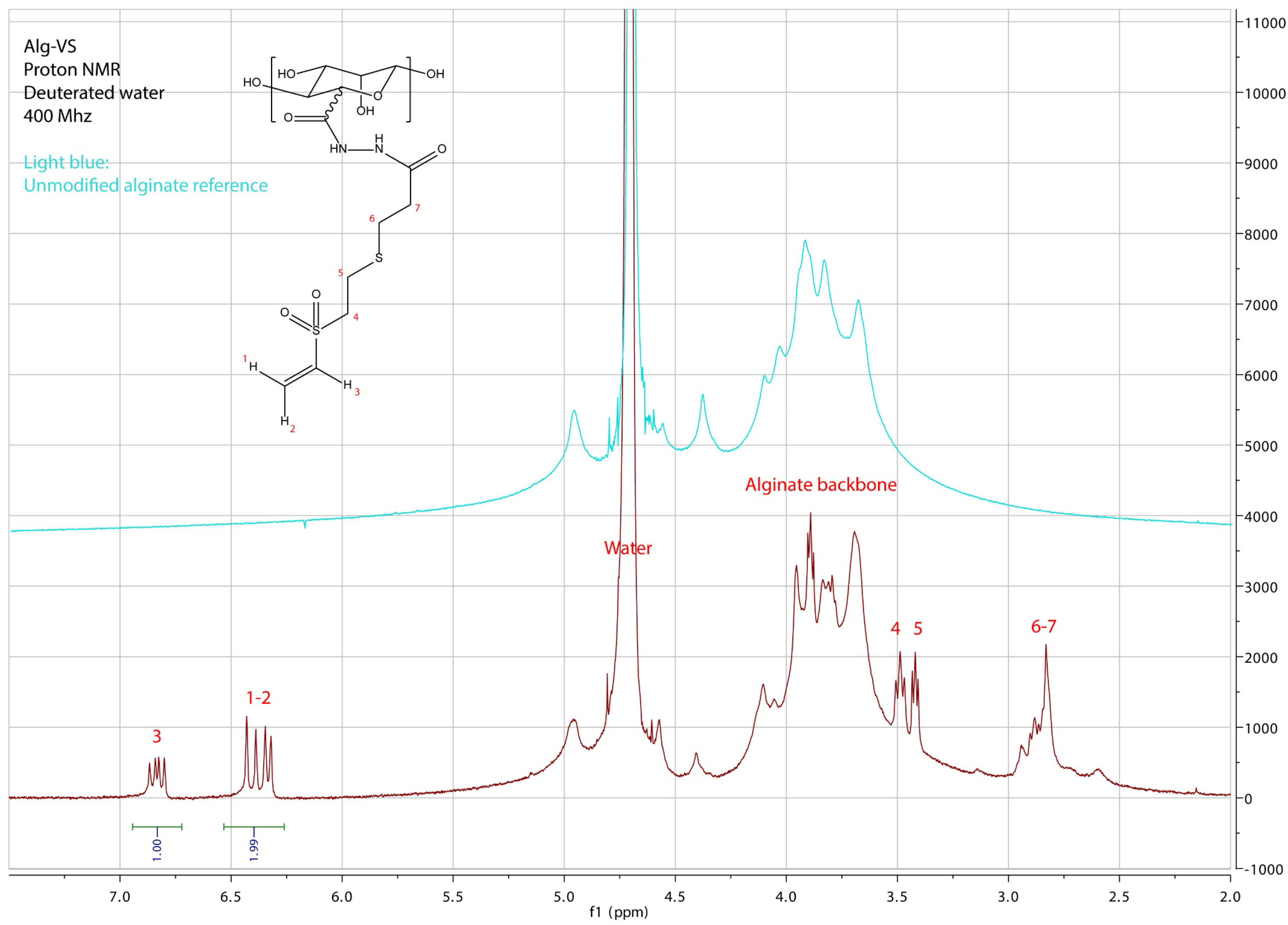

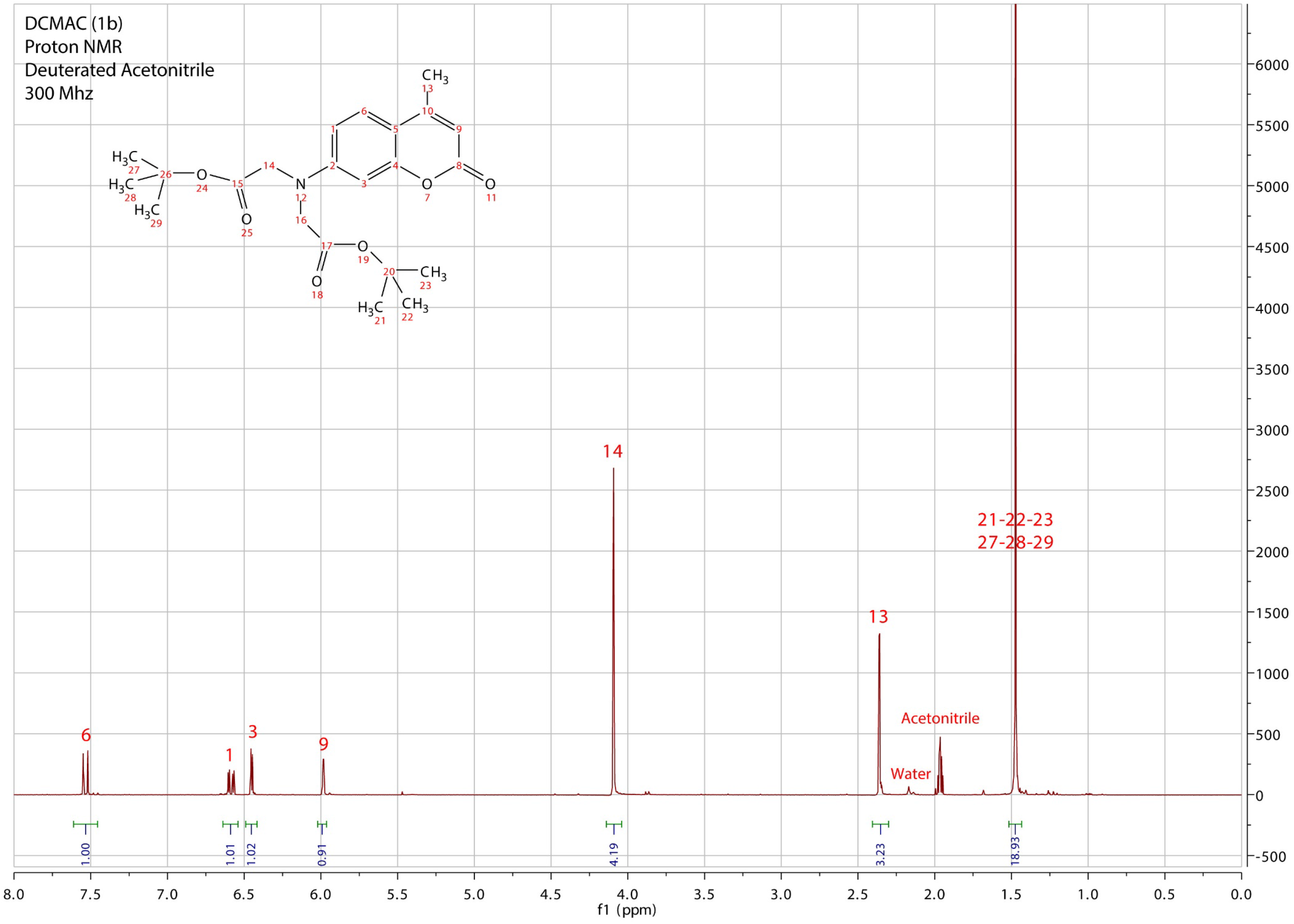

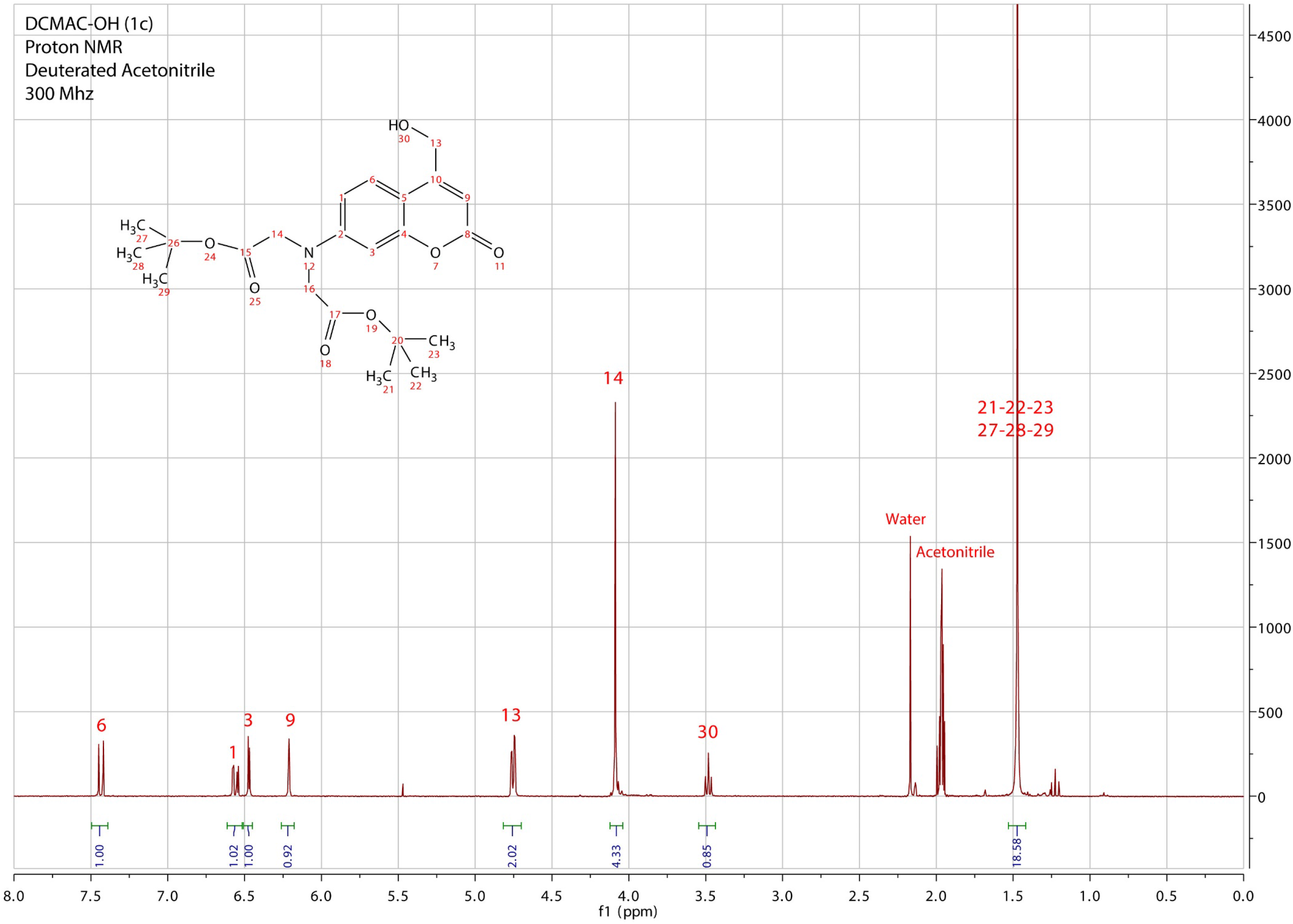

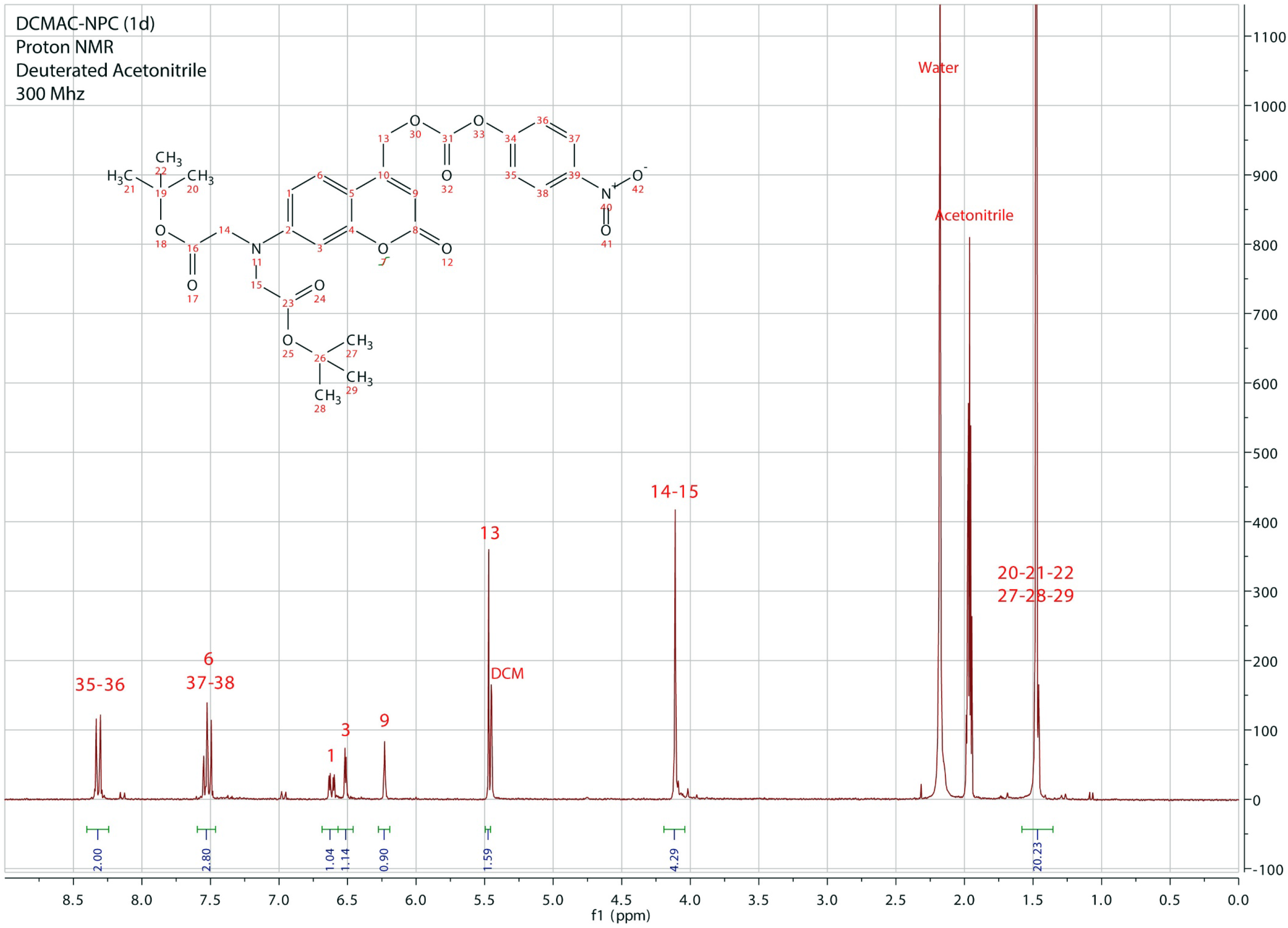

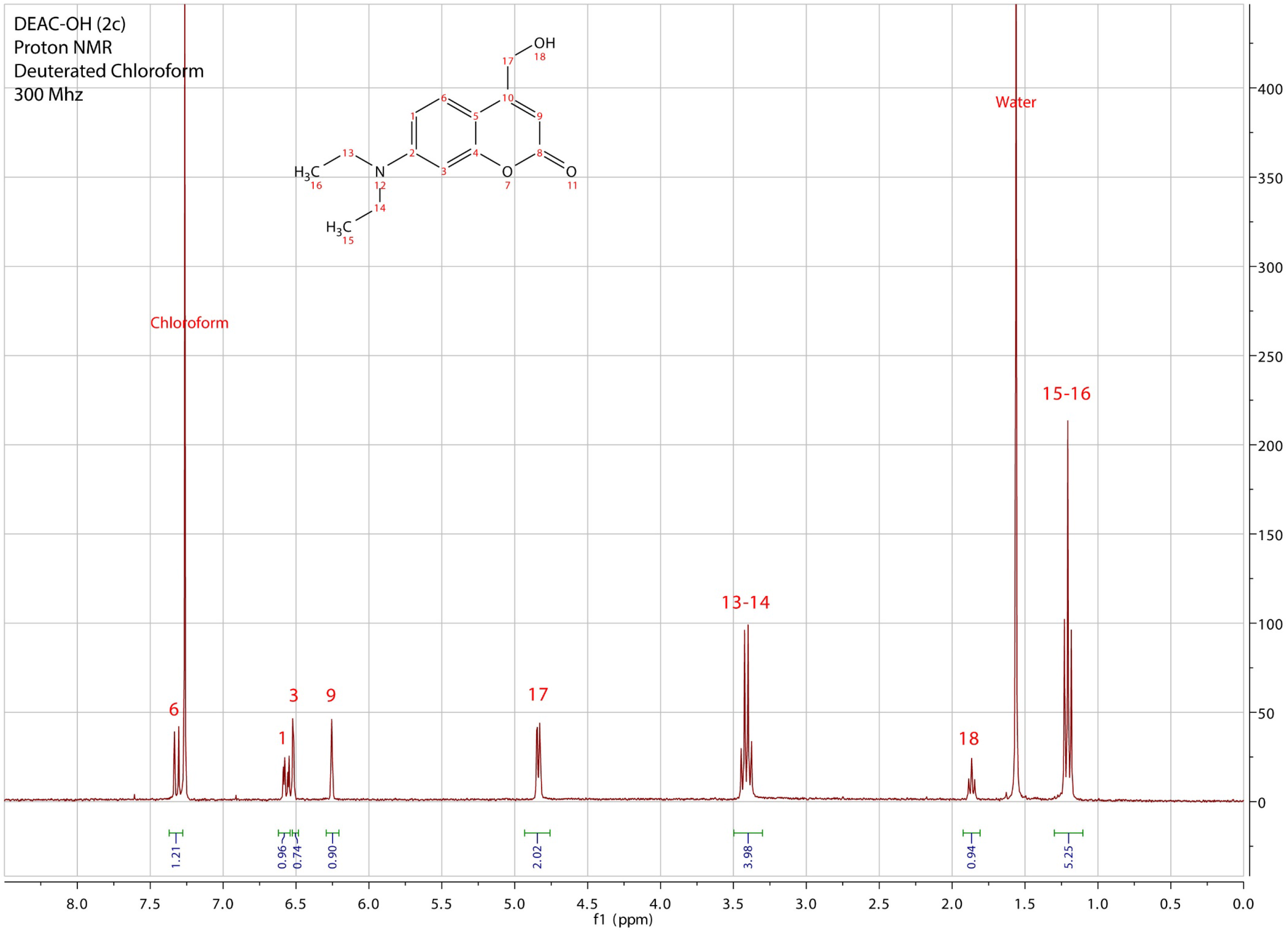

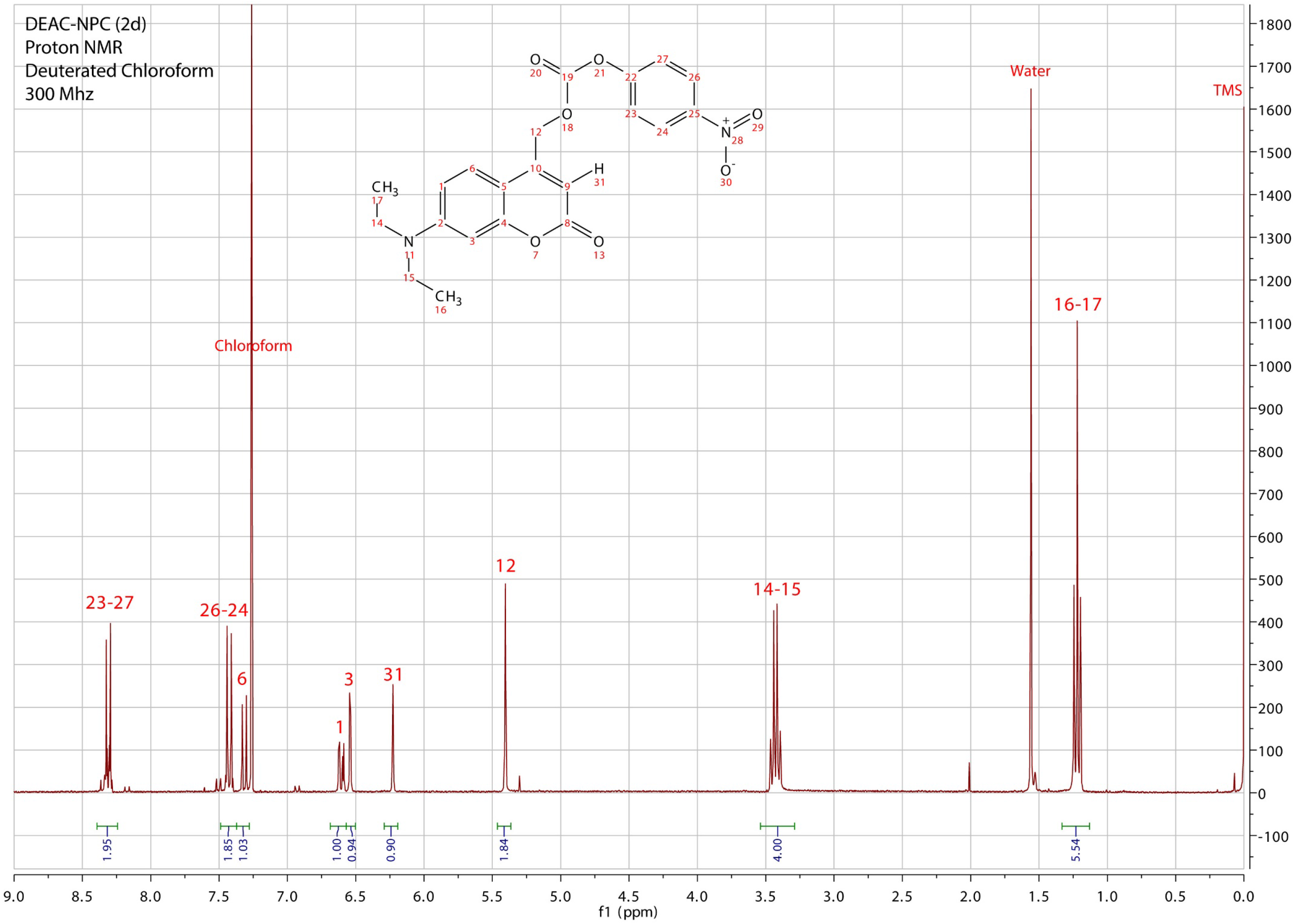

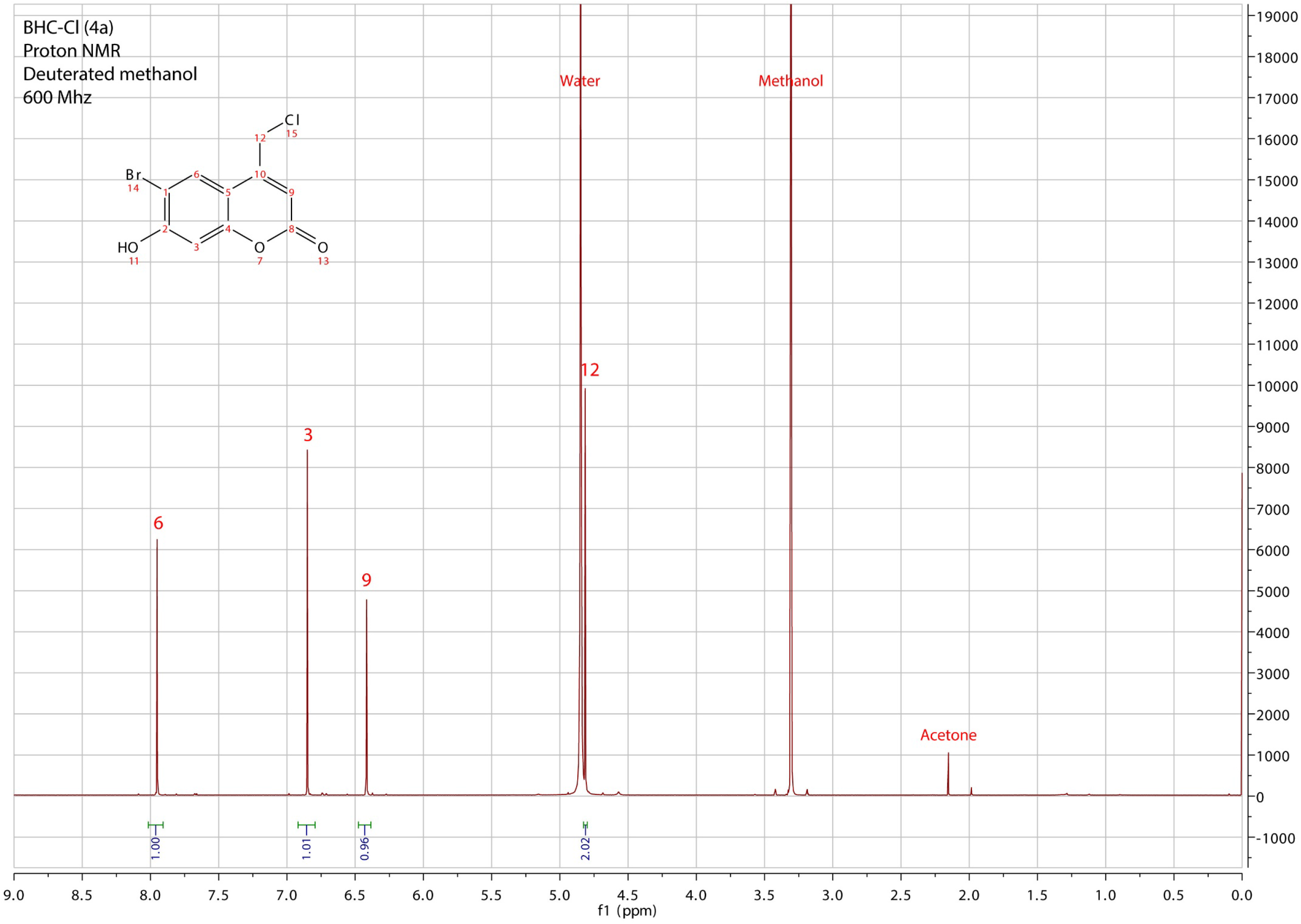

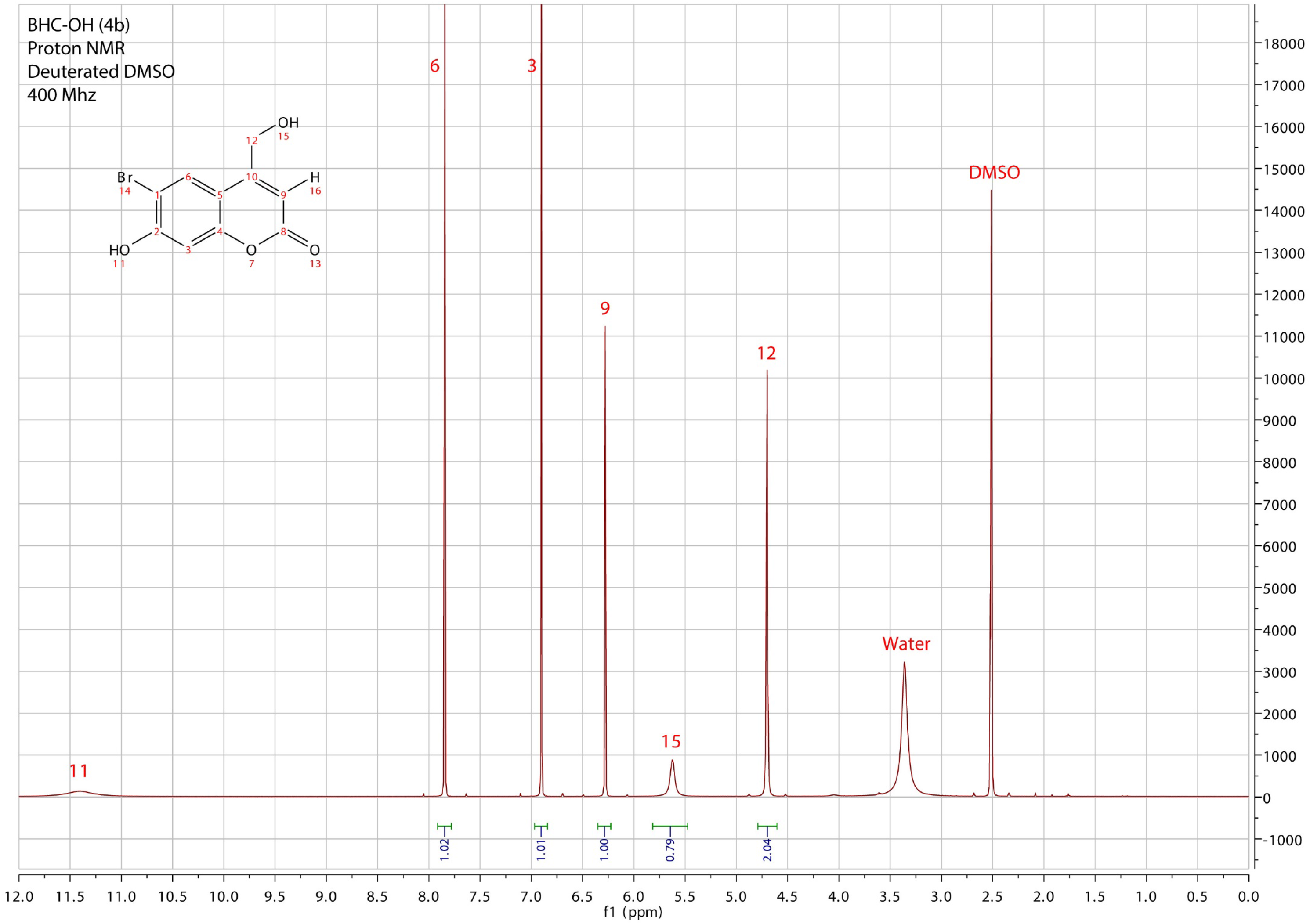

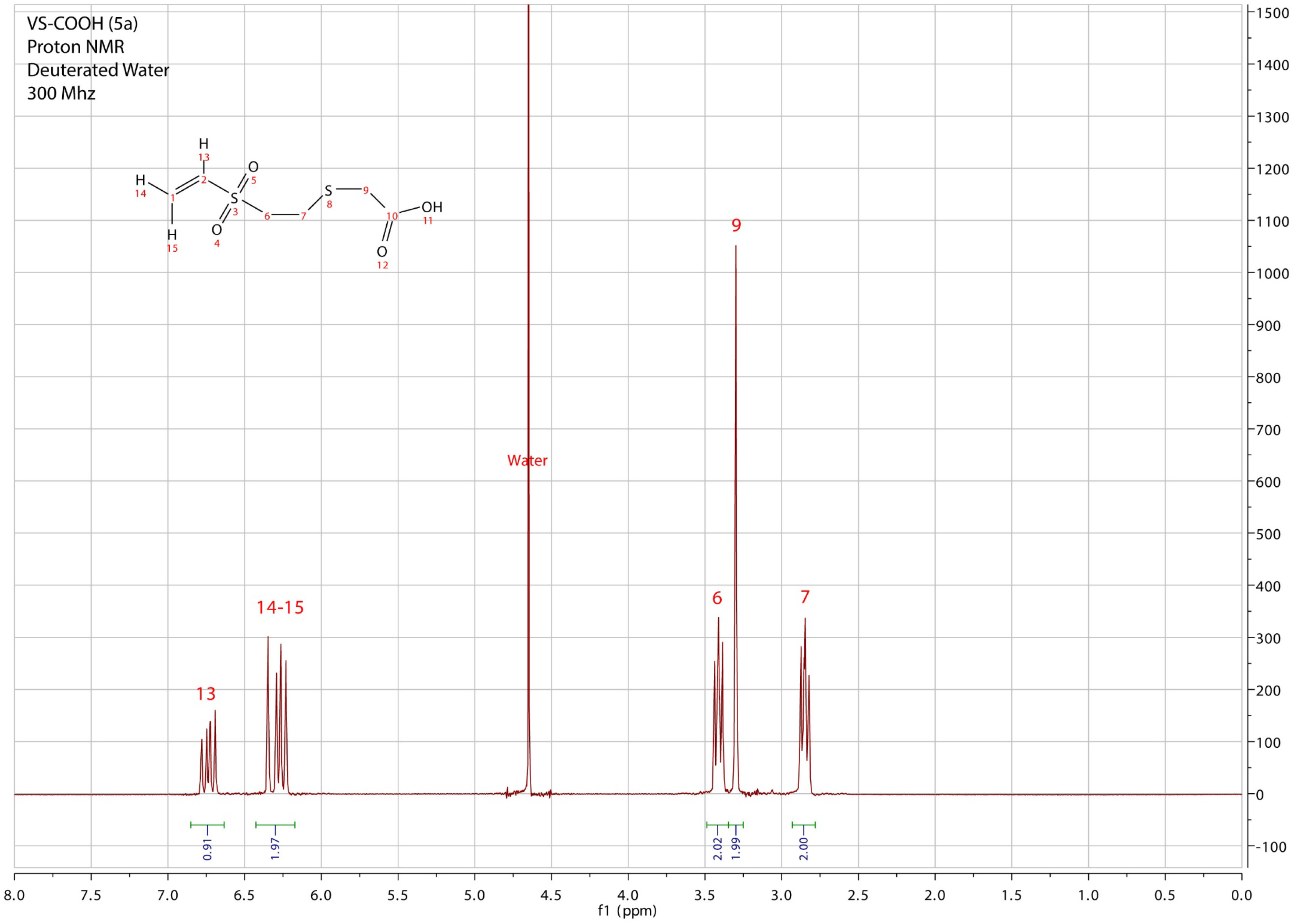

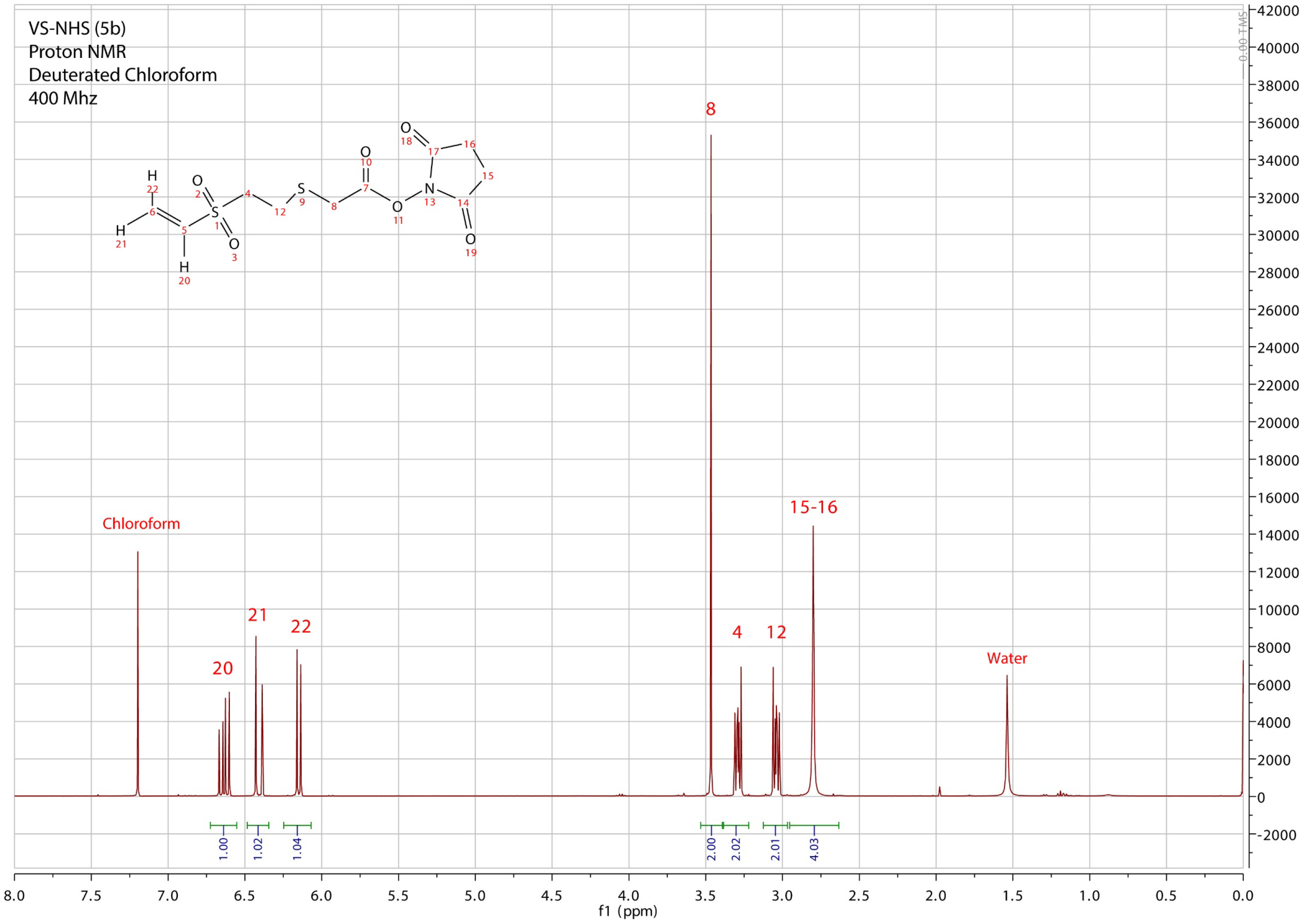

## References

1. Chen, T.-W., Wardill, T. J., Sun, Y., Pulver, S. R., Renninger, S. L., Baohan, A., Schreiter, E. R., Kerr, R. a, Orger, M. B., Jayaraman, V., Looger, L. L., Svoboda, K. & Kim, D. S. Ultrasensitive fluorescent proteins for imaging neuronal activity. Nature 499, 295–300 (2013).

2. Yang, Y., Liu, N., He, Y., Liu, Y., Ge, L., Zou, L., Song, S., Xiong, W. & Liu, X. Improved calcium sensor GCaMP-X overcomes the calcium channel perturbations induced by the calmodulin in GCaMP. Nat. Commun. 9, 1504 (2018).

3. Boyden, E. S., Zhang, F., Bamberg, E., Nagel, G. & Deisseroth, K. Millisecond-timescale, genetically targeted optical control of neural activity. Nat. Neurosci. 8, 1263–1268 (2005).

4. Kandler, K., Katz, L. C. & Kauer, J. A. Focal photolysis of caged glutamate produces long-term depression of hippocampal glutamate receptors. Nat. Neurosci. 1, 119–123 (1998).

5. Wang, X., Chen, X. & Yang, Y. Spatiotemporal control of gene expression by a light-switchable transgene system. Nat. Methods 9, 266–269 (2012).

6. Rost, B. R., Schneider-Warme, F., Schmitz, D. & Hegemann, P. Optogenetic Tools for Subcellular Applications in Neuroscience. Neuron 96, 572–603 (2017).

7. Chou, C., Young, D. D. & Deiters, A. A light-activated DNA polymerase. Angew. Chemie 6064–6067 (2009). doi:10.1002/ange.200901115

8. Deiters, A., Groff, D., Ryu, Y., Xie, J. & Schultz, P. G. A genetically encoded photocaged tyrosine. Angew. Chemie 2794–2797 (2006). doi:10.1002/ange.200600264

9. Dougherty, D. A. Unnatural amino acids as probes of protein structure and function. Curr. Opin. Chem. Biol. 4, 645–652 (2000).

10. Wylie, R. G. & Shoichet, M. S. Three-dimensional spatial patterning of proteins in hydrogels. Biomacromolecules 12, 3789–96 (2011).

11. Lutolf, M. P. Biomaterials: Spotlight on hydrogels. Nat. Mater. 8, 451–3 (2009).

12. Torgersen, J., Qin, X.-H., Li, Z., Ovsianikov, A., Liska, R. & Stampfl, J. Hydrogels for Two-Photon Polymerization: A Toolbox for Mimicking the Extracellular Matrix. Adv. Funct. Mater. n/a-n/a (2013). doi:10.1002/adfm.201203880

13. Kloxin, A. M., Tibbitt, M. W. & Anseth, K. S. Synthesis of photodegradable hydrogels as dynamically tunable cell culture platforms. Nat. Protoc. 5, 1867–1887 (2010).

14. McKinnon, D. D., Brown, T. E., Kyburz, K. a., Kiyotake, E. & Anseth, K. S. Design and characterization of a synthetically accessible, photodegradable hydrogel for user-directed formation of neural networks. Biomacromolecules 15, 2808–2816 (2014).

15. Kloxin, A. M., Kasko, A. M., Salinas, C. N. & Anseth, K. S. Photodegradable hydrogels for dynamic tuning of physical and chemical properties. Science (80-.). 324, 59–63 (2009).

16. Burdick, J. A. & Murphy, W. L. Moving from static to dynamic complexity in hydrogel design. Nat. Commun. 3, 1–8 (2012).

17. Wylie, R. G., Ahsan, S., Aizawa, Y., Maxwell, K. L., Morshead, C. M. & Shoichet, M. S. Spatially controlled simultaneous patterning of multiple growth factors in three-dimensional hydrogels. Nat. Mater. 10, 799–806 (2011).

18. DeForest, C. a, Polizzotti, B. D. & Anseth, K. S. Sequential click reactions for synthesizing and patterning three-dimensional cell microenvironments. Nat. Mater. 8, 659–64 (2009).

19. Li, Z., Ajami, A., Stankevičius, E., Husinsky, W., Račiukaitis, G., Stampfl, J., Liska, R. & Ovsianikov, A. 3D photografting with aromatic azides: A comparison between three-photon and two-photon case. Opt. Mater. (Amst). 35, 1846–1851 (2013).

20. Mosiewicz, K. a, Kolb, L., van der Vlies, A. J., Martino, M. M., Lienemann, P. S., Hubbell, J. a, Ehrbar, M. & Lutolf, M. P. In situ cell manipulation through enzymatic hydrogel photopatterning. Nat. Mater. 12, 1072–8 (2013).

21. Shadish, J. A., Benuska, G. M. & DeForest, C. A. Bioactive site-specifically modified proteins for 4D patterning of gel biomaterials. Nat. Mater. 1 (2019). doi:10.1038/s41563-019-0367-7

22. Strijbis, K., Spooner, E. & Ploegh, H. L. Protein Ligation in Living Cells Using Sortase. 780–789 (2012). doi:10.1111/j.1600-0854.2012.01345.x

23. Pasqual, G., Chudnovskiy, A., Tas, J. M. J., Agudelo, M., Schweitzer, L. D., Cui, A., Hacohen, N. & Victora, G. D. Monitoring T cell-dendritic cell interactions in vivo by intercellular enzymatic labelling. Nat. Publ. Gr. 553, 496–500 (2018).

24. Theile, C. S., Witte, M. D., Blom, A. E. M., Kundrat, L., Ploegh, H. L. & Guimaraes, C. P. Site-specific N-terminal labeling of proteins using sortase-mediated reactions. Nat. Protoc. 8, 1800–1807 (2013).

25. Guimaraes, C. P., Witte, M. D., Theile, C. S., Bozkurt, G., Kundrat, L., Blom, A. E. M. & Ploegh, H. L. Site-specific C-terminal and internal loop labeling of proteins using sortase-mediated reactions. Nat. Protoc. 8, 1787–1799 (2013).

26. Foote, R. S., Cornwell, P., Isham, K. R., Gigerich, H., Stengele, K.-P., Pfleiderer, W. & Sachleben, Ri. A. Photolabile Protecting Groups for Nucleosides:synthesis and Photodeprotection Rates. Tetrahedron 53, 4247–4264 (1997).

27. Bhushan, K. R., DeLisi, C. & Laursen, R. A. Synthesis of photolabile 2-(2-nitrophenyl)propyloxycarbonyl protected amino acids. Tetrahedron Lett. 44, 8585–8588 (2003).

28. Pirrung, M. C., Dore, T. M., Zhu, Y. & Rana, V. S. Sensitized two-photon photochemical deprotection. Chem. Commun. 46, 5313–5315 (2010).

29. Furuta, T., Wang, S. S., Dantzker, J. L., Dore, T. M., Bybee, W. J., Callaway, E. M., Denk, W. & Tsien, R. Y. Brominated 7-hydroxycoumarin-4-ylmethyls: photolabile protecting groups with biologically useful cross-sections for two photon photolysis. Proc. Natl. Acad. Sci. U. S. A. 96, 1193–200 (1999).

30. Wosnick, J. H. & Shoichet, M. S. Three-dimensional Chemical Patterning of Transparent Hydrogels. Chem. Mater. 20, 55–60 (2008).

31. Schönleber, R. O., Bendig, J., Hagen, V. & Giese, B. Rapid Photolytic Release of Cytidine 5 0 -Diphosphate from a Coumarin Derivative : a New Tool for the Investigation of Ribonucleotide Reductases. Bioorg. Med. Chem. 10, 97–101 (2002).

32. Hagen, V., Bendig, J., Frings, S., Eckardt, T., Helm, S., Reuter, D. & Kaupp, U. B. Highly efficient and Ultrafast Phottriggers for cAMP and cGMP by Using Long-Wavelength UV/Vis-Activation. Angew. Chemie - Int. Ed. 40, 1045–1048 (2001).

33. Hagen, V., Dekowski, B., Nache, V., Schmidt, R., Geißler, D., Lorenz, D., Eichhorst, J., Keller, S., Kaneko, H., Benndorf, K. & Wiesner, B. Coumarinylmethyl esters for ultrafast release of high concentrations of cyclic nucleotides upon one- and two-photon photolysis. Angew. Chemie - Int. Ed. 44, 7887–7891 (2005).

34. Hagen, V., Dekowski, B., Kotzur, N., Lechler, R., Wiesner, B., Briand, B. & Beyermann, M. {7-[Bis(carboxymethyl)amino]coumarin-4-yl}methoxycarbonyl derivatives for photorelease of carboxylic acids, alcohols/phenols, thioalcohols/thiophenols, and amines. Chem. - A Eur. J. 14, 1621–1627 (2008).

35. Lyon, R. P., Setter, J. R., Bovee, T. D., Doronina, S. O., Hunter, J. H., Anderson, M. E., Balasubramanian, C. L., Duniho, S. M., Leiske, C. I., Li, F. & Senter, P. D. Self-hydrolyzing maleimides improve the stability and pharmacological properties of antibody-drug conjugates. Nat. Biotechnol. 32, 1059–62 (2014).

36. Mokotoff, M., Mocarski, Y. M., Gentsch, B. L., Miller, M., Zhou, J.-H., Chen, J. & Ball, E. D. Caution in the use of 2-iminothiolane (Traut’s reagent) as a cross-linking agent for peptides. The formation of N-peptidyl-2-iminothiolanes with bombesin (BN) antagonist (d-Trp6,Leu13-psi[CH2NH]-Phe14)BN6-14 and d-Trp-Gln-Trp-NH2. J. Pept. Res. 57, 383–389 (2001).

37. Morpurgo, M., Veronese, F. M., Kachensky, D. & Harris, J. M. Preparation of characterization of poly(ethylene glycol) vinyl sulfone. Bioconjug. Chem. 7, 363–8 (1996).

38. Ostuni, E., Chapman, R. G., Holmlin, R. E., Takayama, S. & Whitesides, G. M. A survey of structure-property relationships of surfaces that resist the adsorption of protein. Langmuir 17, 5605–5620 (2001).

39. Broguiere, N., Isenmann, L. & Zenobi-Wong, M. Novel enzymatically cross-linked hyaluronan hydrogels support the formation of 3D neuronal networks. Biomaterials 99, 47–55 (2016).

40. Deforest, C. A. & Anseth, K. S. Cytocompatible click-based hydrogels with dynamically tunable properties through orthogonal photoconjugation and photocleavage reactions. Nat. Chem. 3, 925–931 (2011).

41. Fisher, S. A., Tam, R. Y., Fokina, A., Mahmoodi, M. M., Distefano, M. D. & Shoichet, M. S. Photo-immobilized EGF chemical gradients differentially impact breast cancer cell invasion and drug response in defined 3D hydrogels. Biomaterials (2018). doi:10.1016/j.biomaterials.2018.01.032

42. Skylar-Scott, M. A., Liu, M. C., Wu, Y., Dixit, A. & Yanik, M. F. Guided Homing of Cells in Multi-Photon Microfabricated Bioscaffolds. Adv. Healthc. Mater. 5, 1233–1243 (2016).

43. Ovsianikov, A., Li, Z., Torgersen, J., Stampfl, J. & Liska, R. Selective Functionalization of 3D Matrices Via Multiphoton Grafting and Subsequent Click Chemistry. Adv. Funct. Mater. n/a-n/a (2012). doi:10.1002/adfm.201200419

44. Levi-Montalcini, R. The nerve growth factor 35 years later. Science (80-.). 237, 1154–1162 (1987).

45. Sarig-Nadir, O. & Seliktar, D. Compositional alterations of fibrin-based materials for regulating in vitro neural outgrowth. Tissue Eng. Part A 14, 401–11 (2008).

46. Schense, J. C., Bloch, J., Aebischer, P. & Hubbell, J. a. Enzymatic incorporation of bioactive peptides into fibrin matrices enhances neurite extension. Nat. Biotechnol. 18, 415–9 (2000).

47. Chan, C. C. M., Roberts, C. R., Steeves, J. D. & Tetzlaff, W. Aggrecan components differentially modulate nerve growth factor–responsive and neurotrophin-3-responsive dorsal root ganglion neurite growth. J. Neurosci. Res. 86, 581–592 (2008).

48. Sakiyama-elbert, S. E., Panitch, A. & Hubbell, J. A. Development of growth factor fusion proteins for cell-triggered drug delivery. FASEB J. (2001).

49. Sakiyama-Elbert, S. E. & Hubbell, J. a. Controlled release of nerve growth factor from a heparin-containing fibrin-based cell ingrowth matrix. J. Control. Release 69, 149–58 (2000).

50. Broguiere, N., Cavalli, E., Salzmann, G. M., Applegate, L. A. & Zenobi-Wong, M. Factor XIII Cross-Linked Hyaluronan Hydrogels for Cartilage Tissue Engineering. ACS Biomater. Sci. Eng. 2, 2176–2184 (2016).

